# The C-terminal tail guides assembly and degradation of membrane proteins

**DOI:** 10.1101/843326

**Authors:** Sha Sun, Malaiyalam Mariappan

**Affiliations:** Department of Cell Biology, Nanobiology Institute, Yale School of Medicine, Yale West Campus, West Haven, CT 06516, USA

## Abstract

A large number of newly synthesized membrane proteins in the endoplasmic reticulum (ER) are assembled into multi-protein complexes, but little is known about the mechanisms required for either assembly or degradation of unassembled membrane proteins. We find that C-terminal transmembrane domains (C-TMDs) with shorter tails are inefficiently inserted into the ER membrane since the translation is terminated before they emerge from ribosomes. These TMDs of insufficient hydrophobicity are post-translationally retained by the Sec61 translocon, thus providing a time window for efficient assembly with TMDs from partner membrane proteins. The unassembled C-TMDs are slowly flipped into the ER lumen. While the luminal chaperone BiP captures flipped C-TMDs with long tails and routes them to the ER-associated quality control, C-TMDs with shorter tails are diffused into the nuclear membrane. Thus, our studies suggest that C-terminal tails harbor important biochemical features for both biosynthesis and quality control of membrane protein complexes.

## INTRODUCTION

Membrane proteins represent one-third of the proteins encoded by the human genome and carry out essential cellular processes including molecular transport, signaling, and cell-cell communication (Krogh et al., 2001; Shao and Hegde, 2011). Most membrane proteins utilize the co-translational pathway for insertion into the endoplasmic reticulum (ER). As the hydrophobic signal sequence or the first transmembrane domain (TMD) of a membrane protein emerges from the cytosolic ribosome, the signal recognition particle (SRP) captures it and delivers the ribosome nascent chain complex to the ER membrane via the SRP receptor (Grudnik et al., 2009; Park and Rapoport, 2012; Zhang and Shan, 2014). The SRP bound TMD is then transferred to the Sec61 translocon channel in the ER membrane. The hydrophobic TMD of a nascent membrane protein engages the lateral gate of the channel, where it can sample the hydrophobic chains of phospholipids. Subsequently, the TMD partitions into the lipid phase as the nascent chain further elongates during translation. It is generally believed that TMDs of membrane proteins are sequentially inserted into the ER membrane (Skach, 2007).

It is estimated that over 40% of newly synthesized membrane proteins assemble into multi-protein complexes (Juszkiewicz and Hegde, 2018). Even though these multi-membrane protein complexes serve fundamental functions in cells, the mechanisms that govern the assembly of individual membrane proteins into complexes are poorly understood. The assembly of newly synthesized membrane proteins faces several challenges in the two-dimensional ER membrane. For instance, unassembled membrane proteins can have non-productive interactions, leading to misfolding, aggregation and degradation(Balchin et al., 2016; Kramer et al., 2019). Second, since the ER is the largest membrane network within eukaryotic cells, the newly synthesized subunits can move from the rough ER to the other part of the ER (Foresti et al., 2014; Khmelinskii et al., 2014; Wu et al., 2018), thus diluting their concentration as well as reducing their chance to find each other. Third, proteins have often the propensity to form homo-oligomers since they are translated from the same polysome and their local subunit concentration will be high, thus reducing the chance to assemble with the partner protein. In recent years, much attention has been focused on understanding the assembly of soluble protein complexes (Schwarz and Beck, 2019). These studies propose two prevailing models. First, the large protein complexes such as proteasome often have dedicated molecular chaperones that can shield the nascent proteins from inappropriate interactions and facilitate assembly with partner proteins (Le Tallec et al., 2007). Second, the newly synthesized proteins can be co-translationally assembled into hetero-oligomeric complexes in eukaryotes (Shiber et al., 2018).

In contrast to soluble protein complexes, less is known about the assembly of membrane proteins complexes in the ER membrane, where the majority of the membrane proteins are synthesized in cells. Pioneering early studies have used T-cell receptor (TCR) complex as a model membrane protein complex to investigate this problem (Manolios et al., 1990). The opposite charge residues in TMDs of membrane proteins drive assembly by forming ionic-bonds between subunits (Bonifacino et al., 1990; Cosson et al., 1991; Manolios et al., 1990). The unassembled TCR subunits can be translocated completely into the ER lumen due to their marginally hydrophobic TMDs (Feige and Hendershot, 2013). However, it is unclear how these two different subunits find each other in the extensive ER membrane network after their synthesis and form a specific ionic pair. Moreover, this is even less understood for assembly of polytopic membrane protein complexes, but it is often proposed that membrane chaperones may mediate the assembly process. However, experimental evidence of membrane chaperones mediating the assembly of membrane protein complexes is lacking.

Recent proteomic based pulse chase experiments suggest that many newly synthesized multi-subunit membrane proteins fail to assemble with their partner proteins (McShane et al., 2016). It is unclear how these orphaned membrane subunits are selectively recognized by ER-associated degradation (ERAD) pathways (Brodsky, 2012). Early studies have extensively used two single spanning membrane proteins, TCRα and CD3δ, as models for the degradation of unassembled proteins by ERAD (Bonifacino and Lippincott-Schwartz, 1991; Bonifacino et al., 1989; Huppa and Ploegh, 1997). Two models have been proposed on how unassembled membrane proteins are recognized by ERAD. In the first model, unassembled membrane proteins may expose polar residues, which are normally used for the assembly with the partner protein, within the lipid bilayer. These polar residues in TMDs may be recognized by ERAD components for degradation(Kikkert et al., 2004; Sato et al., 2009). However, the direct evidence for recognition of polar residues in TMDs of orphaned proteins by ERAD machinery is unclear (Ruggiano et al., 2014). In the second model, the presence of polar or charged residues in TMDs renders them to translocate into the ER lumen (Feige and Hendershot, 2013; Shin et al., 1993). These translocated proteins are then efficiently recognized by ERAD (Feige and Hendershot, 2013). It remains to be determined whether this model applies to all orphaned membrane proteins, especially for polytopic membrane proteins (Juszkiewicz and Hegde, 2018).

To reveal the biochemical features that are necessary for both assembly and degradation of membrane protein complexes, we chose to investigate the assembly of the tail-anchored membrane protein insertion complex composed of WRB and CAML proteins (Vilardi et al., 2011; Vilardi et al., 2014; Yamamoto and Sakisaka, 2012). Our studies suggest that the newly synthesized membrane proteins are prone to be diffused into other parts of the ER such as nuclear membrane, thus cells have evolved with mechanisms to limit the diffusion to improve the assembly efficiency. We identify that the Sec61 translocon can post-translationally retain newly synthesized membrane proteins. This retention mechanism depends on less hydrophobic C-TMDs with shorter tails. These TMDs are neither efficiently inserted or translocated by the Sec61 translocon because the translation is terminated before they emerge from ribosomes. These Sec61 translocon trapped TMDs can efficiently assemble with TMDs from partner membrane proteins. If they missed assembly with their partner TMDs, C-TMDs are slowly flipped into the ER lumen. While C-TMDs with short tails are diffused into the nuclear membrane, the luminal BiP captures the C-TMDs with long tails and route them to the Hrd1 E3 ligase complex for degradation. Thus, our studies suggest that the C-terminal cytosolic tails contain crucial signals for both assembly and quality control of membrane protein complexes.

## RESULTS

### Assembly failed membrane proteins are degraded by the ubiquitin-proteasome pathway

To determine the biochemical features that are required for the assembly of multi-membrane protein complexes, we sought for model membrane protein substrates that should fulfill three criteria. First, they must be relatively small and amenable for biochemical manipulations. Second, they should be quickly degraded when they failed to assemble. Third, degradation signals or degrons should be buried when they are assembled, but exposed when they fail to assemble. We reasoned that tail-anchored membrane protein insertase complex, WRB and CAML, may be ideal substrates to investigate this fundamental problem since both proteins are relatively small in their sizes (20 and 33 kDa, respectively) with each having three TMDs (Figure 1A, Figure S1A). We first tested whether these proteins are degraded when transiently expressed individually in HEK293 cells. The cycloheximide chase assay revealed that the expression of either WRB or CAML alone led to degradation in a proteasomal dependent manner since the turnover could be inhibited by the proteasomal inhibitor MG132 (Figure 1B, C). However, the co-expression of both WRB and CAML slowed their degradation compared to when they were expressed separately (Figure 1D). We surmised that the coexpression of WRB with CAML did not completely stabilize WRB and CAML because of the inefficient assembly during their synthesis in the ER. To test this, we made a fusion construct of WRB and CAML based on the topology of their yeast counterparts as previously described (Wang et al., 2014). The yeast homologous fusion protein Get2-Get1 protein is functional for TA protein insertion, suggesting that the predicted topology of CAML-WRB is likely correct (Figure 1A). The expression of the CAML-WRB construct was very stable as no significant degradation was observed within 2h of the chase period. (Figure 1E). To confirm that the degradation of WRB and CAML are ubiquitin-dependent degradation by the proteasome, we immunoprecipitated the proteins from cells and immunoblotted them to detect HA-tagged polyubiquitin chains. A strong smear of polyubiquitinated bands was observed when WRB or CAML expressed alone compared to when they were expressed together or the fusion protein (Figure 1F). To further determine the unassembled WRB and CAML were retrotranslocated as full-length proteins into the cytosol, we isolated the cytosol fraction after cells were treated with MG132. The full-length WRB and CAML could be detected in the cytosol when they were expressed separately (Figure 1G). In contrast, CAML-WRB fusion or coexpression of both showed a little signal in the cytosol, indicating that the assembled complex is not efficiently recognized by ERAD machinery. The cells treated with a combination of both MG132 and an inhibitor of p97 ATPase (Magnaghi et al., 2013), which is essential to extract misfolded proteins from the membrane, served as negative controls because unassembled WRB and CAML were not detected in cytosol fractions from these cells.

**Figure 1.**
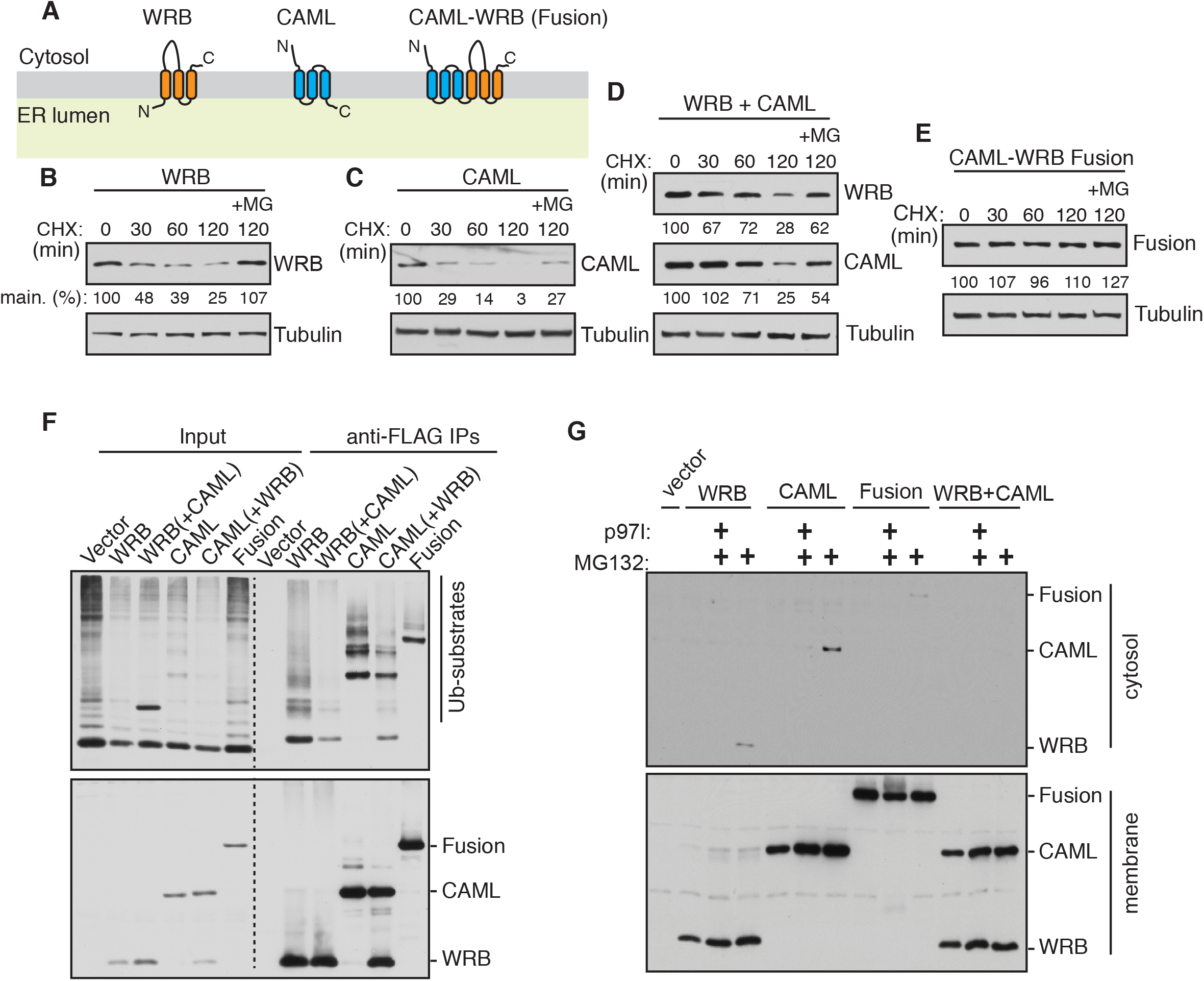
Unassembled membrane proteins are degraded by the ubiquitin-proteasome pathway. (A) Schematics showing the predicted topologies of WRB, CAML and CAML-WRB fusion. The fusion protein is composed of CAML and WRB with a 15 amino acid linker in between. (B) WRB-FLAG was transfected into HEK293 cells. After 24h of transfection, cells were treated with cycloheximide (CHX) for the indicated time points and analyzed by immunoblotting with an anti-FLAG antibody for WRB. Tubulin was immunoblotted as a loading control. The protein level at 0-hour time point was taken as 100%, and the percentage of the remaining protein was calculated with respect to 0 hour. (C)-(E) The cycloheximide chase experiments were performed and analyzed for the indicated constructs as in (B). (F) HEK293 cells transiently expressing the indicated constructs with Ubiquitin-HA were immunoprecipitated with anti-FLAG beads and analyzed by immunoblotting with anti-HA antibodies for ubiquitinated substrates and an anti-FLAG antibody for CAML and WRB. (G) HEK293 cells expressing indicated constructs were treated with MG132 alone or MG132 with an P97 ATPase inhibitor (P97I). The cells were permeabilized with digitonin to collect cytosol and membrane fractions. The samples were analyzed by immunoblotting for the indicated antigens using an anti-FLAG antibody. Note that the retrotranslocated WRB or CAML appears in the cytosol when cells were treated with the proteasome inhibitor MG132, but not when treated with both P97 inhibitor and MG132.

### C-TMDs with insufficient hydrophobicity serve as degrons when flipped into the ER lumen

We hypothesized that the unassembled membrane proteins must expose a degradation signal or degron for recognition by ERAD. To investigate this, we focused on WRB as a model substrate. We reasoned that the degron must lie in one of three TMDs of WRB since WRB lacks a prominent luminal domain. Also, the cytosolic tryptophan-rich domain of WRB is functional without the rest of the protein (Vilardi et al., 2011). Therefore, we swapped one TMD at a time with a stable TMD from either Sec61β or Ost4p (Figure 2A and Figure S1B) based on the previously described protocol (Wang et al., 2014). Replacing the first TMD of WRB led to the degradation similar to the wild type (Figure 2B). The second TMD swap exhibited a slightly faster degradation. Remarkably, swapping the 3rd TMD resulted in complete stabilization of WRB, suggesting that the 3rd TMD contains a degron (Figure 2B). The close inspection of the amino acid sequence of the C-TMD revealed that it has a single positively charged lysine residue (Figure 2C). To test the lysine residue in the C-TMD is required for the recognition by ERAD, we mutated lysine to hydrophobic leucine residue and analyzed by protein turn over assay. In agreement with our view, the degradation of WRB (K164L) was completely inhibited compared to the wild type (Figure 2D), confirming that the lysine residue in the C-TMD serves as a degron. To our surprise, replacing the lysine residue with either a charged residue or a hydrophilic residue caused destabilization (Figure 2D). These results suggested that the lysine residue in the C-TMD of WRB is not the direct signal for degradation, but rather indicated that the overall hydrophobicity of the C-TMD may play a role in recognition by quality control factors. This led us to ask whether the C-TMD is properly inserted into the ER membrane since it has a relatively low apparent free energy (ΔGapp=+2.28) (Hessa et al., 2005; Hessa et al., 2007). Accordingly, a negative value of free energy indicates that the sequence can be efficiently recognized as a TMD by the Sec61 translocon and integrated into the lipid bilayer. Conversely, a positive value of free energy indicates that the sequence is less efficiently inserted unless it is helped by interactions with neighboring TMDs. To test if the C-TMD of WRB is flipped into the ER lumen, we appended a glycosylation site (NFT) at the C terminus of WRB (Figure 2E and Figure S1C). Interestingly, about 37% of the C-TMD of WRB was flipped into the ER lumen (Figure 2F). The glycosylated band of WRB was verified by the treatment with endoglycosidase H (Endo H). Consistent with WRB without a glycosylation tag, the glycosylated (flipped) form of WRB was also quickly degraded as shown by the pulse-chase experiment (Figure 2G). To verify the predicted topology of CAML with N-terminus in the cytosol and C-terminus in the ER lumen (Figure 1A), we appended an N-glycosylation motif to the C-terminus of CAML. As predicted, the majority of CAML appeared to be correctly oriented in the membrane since its C-terminus was efficiently glycosylated by the ER luminal glycosylation machinery (Figure S2A). Taken together, a lysine in the C-TMD of WRB causes the flipping of the TMD into the ER lumen, thus allowing its capture by quality control machinery.

**Figure 2.**
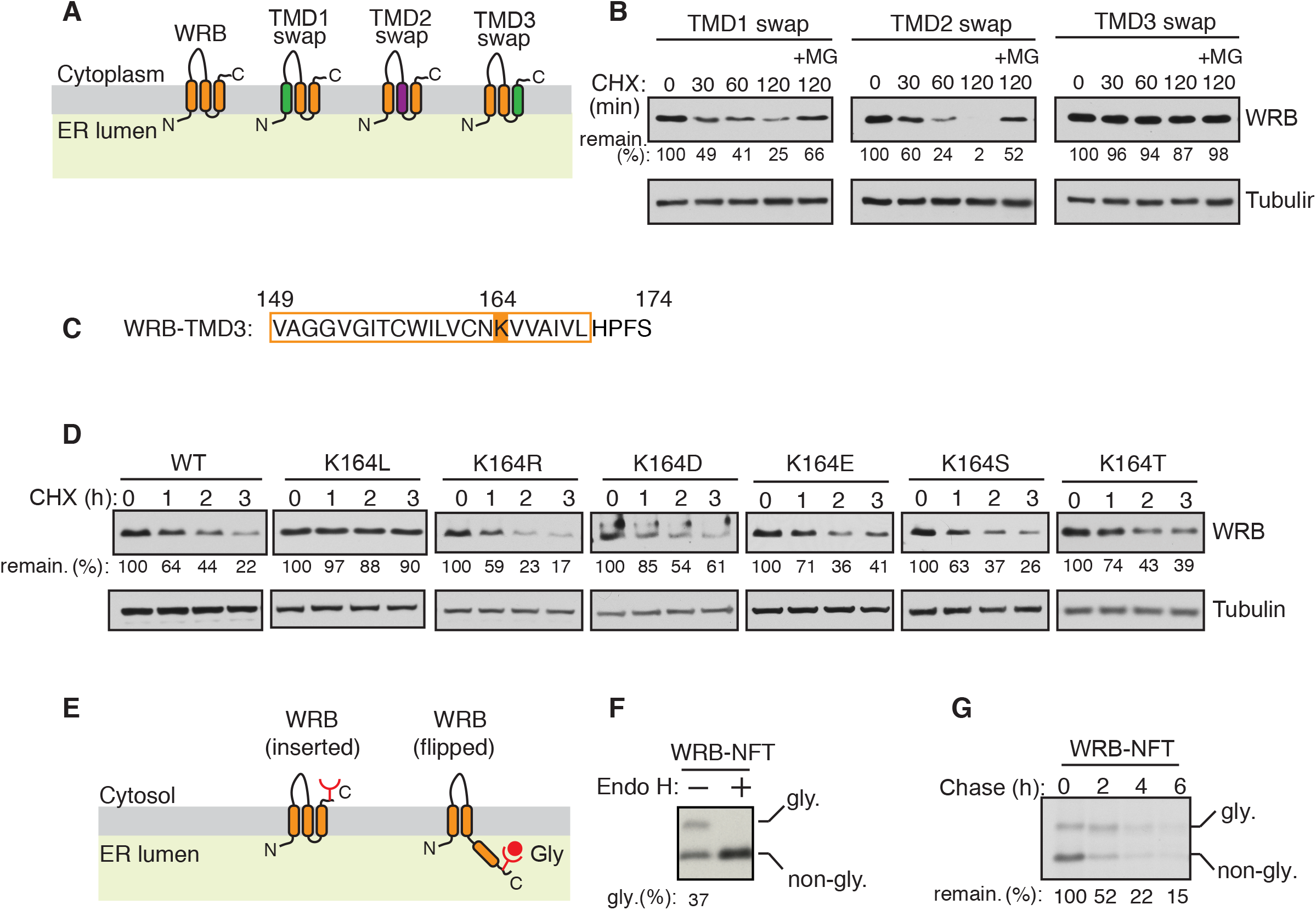
A hydrophilic residue in the C-TMD of WRB serves as a degron by flipping the TMD into the ER lumen. (A) Schematics showing the swapping of transmembrane domains (TMDs) of WRB with either the TMD from Ost4p for TMD1 and TMD3 swaps (shown in green) or the TMD from Sec61β for the TMD2 swap (shown in purple). (B) The indicated WRB swaps were transfected into HEK293 cells. After 24h of transfection, cells were treated with cycloheximide (CHX) for the indicated time points and analyzed by immunoblotting with an anti-FLAG antibody. Tubulin was immunoblotted as a loading control. The protein level at 0-hour time point was taken as 100%, and the percentage of the remaining protein was calculated with respect to 0 hour. (C) A single lysine (K164) is indicated in orange in the amino acid sequence of WRB TMD3. (D) The cycloheximide chase experiments were performed and analyzed for the indicated constructs as in (B). (E) A schematic of two different topologies of WRB (inserted and flipped forms) is shown. The glycosylation acceptor site NFT is added after the C-terminus HA tag of WRB. When the C-TMD is flipped into the ER lumen, the NFT site becomes glycosylated. (F) The lysate from WRB-HA-NFT transfected cells was treated without or with endoglycosidase H (Endo H) and analyzed by immunoblotting with an anti-HA antibody. The percentage of flipped form (glycosylated) from total was quantified with Image J and shown under the blot. (G) WRB-HA-NFT transfected cells were metabolically labeled and chased for the indicated time points and analyzed by autoradiography after immunoprecipitation with anti-HA antibody beads.

We next asked whether the C-TMD flipping into the ER lumen is a common feature for unassembled membrane proteins. To address this, we looked for other membrane protein complexes. The Sec61 translocon is composed of three subunits, α, β, and γ. We reasoned that the C-TMD of α subunit might flip into the ER lumen if it failed to assemble with other two subunits. To test this, we added an N-glycosylation motif to the C-terminus of Sec61α and monitored its flipping into the lumen. Consistent with our prediction, a small fraction of Sec61α was flipped into the ER lumen as shown by an Endo H-sensitive glycosylated band (Figure S3A). We next tested the flipping of the C-TMD of TRAPγ, which forms a heterotrimeric complex with α and β subunits (Lang et al., 2017). Similar to WRB, about 50% of TRAPγ was flipped into the ER lumen as shown by the glycosylation assay (Figure S3B). We further examined whether the flipping of the C-TMD of a polytopic membrane protein might be an indicator of defects in assembly of TMDs within the same protein. To test this, we used peripheral myelin protein 22 (PMP22) as a model substrate since many patients affected by Charcot-Marie-Tooth disease possess mutations in TMDs of PMP22. We reasoned that introducing a patient mutation into the C-TMD of PMP22 (Navon et al., 1996) might flip the TMD into the ER lumen. Consistent with our notion, introducing the positively charged arginine residue in the C-TMD of PMP22 led to its flipping and glycosylation in the ER lumen (Figure S3C). Collectively, these findings suggest that the flipping of the C-TMD is a general phenomenon for unassembled membrane proteins.

### The C-terminal cytosolic tail influences both flipping and proper insertion of C-TMDs

We investigated whether the flipping of C-TMD into the ER lumen is caused by its low hydrophobicity. To test this, we constructed variants of C-TMDs with either increasing hydrophobicity by introducing hydrophobic leucine residues or decreasing hydrophobicity by introducing hydrophilic residues (Figure 3A). We analyzed the flipping of C-TMDs of WRB variants by the glycosylation assay combined with immunoblotting. C-TMDs of WRBs with insufficient hydrophobicity flipped into the ER lumen as evidenced by glycosylated bands (Figure 3B). The efficiency of the C-TMD flipping generally correlated with its hydrophobicity. For instance, about 90% of very hydrophilic C-TMD of WRB (DDRRK) flipped into the ER lumen. Strikingly, increasing the hydrophobicity of C-TMD did not prevent its flipping into the ER lumen, arguing that strong hydrophobicity of a C-TMD is not a determinant for insertion into the membrane (Figure 3B). This result is further corroborated with the data derived from metabolically labeled cells expressing WRB variants as well as the data from an in vitro experiment where WRB variants were translated in the presence of rough microsomes (Figure 3C and Figure S4).

**Figure 3.**
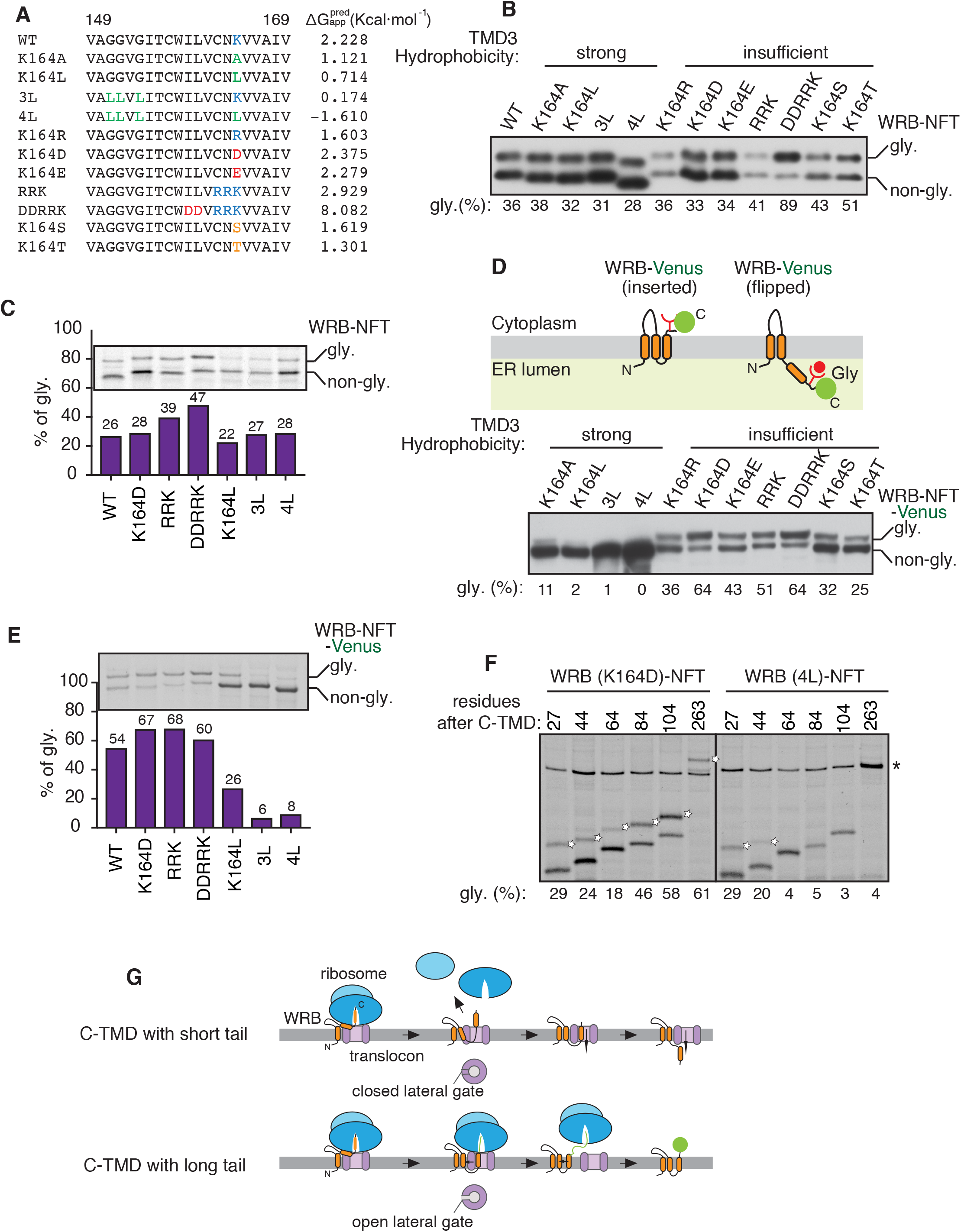
The cytosolic tail influences both insertion and flipping of the C-TMD. (A) Amino acid sequences of WRB TMD3 and its variants along with apparent free energy predictions The mutations are highlighted using the following colours. Hydrophobic residue: Green. Positively charged residues: Blue. Negatively charged residues: Red. Hydrophilic residues: Orange. (B) The indicated WRB-HA-NFT variants were transfected into HEK293 cells and analyzed by immunoblotting with an anti-HA antibody. The percentage of flipped form (glycosylated) was quantified with Image J and shown under the blot. (C) HEK293 cells expressing the indicated WRB-HA-NFT variants were metabolically labelled for 30min and immunoprecipitated with anti-HA antibody beads. The immunoprecipitants were analyzed by autoradiography. The percentage of flipped form (glycosylated) was quantified and plotted in a bar graph (bottom). (D) Top panel: A schematic showing the two different topologies of WRB-NFT-Venus with inserted or flipped C-TMD. The glycosylation acceptor site NFT is added between WRB and Venus. Bottom panel: the indicated WRB-NFT-Venus variants were transfected into HEK293 cells and analyzed as in (B). (E) HEK293 cells expressing the indicated WRB-NFT-Venus variants were analyzed as in (C). (F) The indicated WRB-NFT-Venus constructs with C-terminal truncations were transfected and analyzed as in (C). Empty stars denote flipped forms of WRB variants. Star indicates the non-specific band. (G) A schematic showing the C-terminal tail influencing the insertion of the C-TMD with strong hydrophobicity. Note that the open and close states of the translocon lateral gate are influenced by the ribosome binding.

We asked why the C-TMD with strong hydrophobicity does not obey the biological hydrophobicity, and thus is not efficiently inserted into the membrane. We reasoned that when the translation is terminated, most of the sequence of C-TMD and its tail might still be within the exit tunnel of the ribosome, which can accommodate ~40 amino acids (Voss et al., 2006)(Figure S1C). This might force the translocon to post-translationally recognize and insert the C-TMD into the lipid bilayer. We hypothesized this post-translational recognition of C-TMD by the Sec61 translocon might be slow and inefficient because the previous structural studies have shown that the lateral gate of the translocon is closed in the absence ribosome (Voorhees and Hegde, 2016a; Wu et al., 2019). To test this idea, we increased the length of the cytosolic C-tail by appending a large Venus tag which comprises 238 amino acids (Figure S1D). In support of our hypothesis, the C-TMD flipping of WRB-Venus variants with strong hydrophobicity was significantly prevented as shown by both immunoblotting and metabolic labeling results (Figure 3D, E). Consistent with the biological hydrophobicity scale (Hessa et al., 2007), C-terminal TMDs with insufficient hydrophobicity flipped into the ER lumen. Of note, WRB-Venus with insufficient hydrophobicity exhibited more efficient flipping into the ER lumen compared to WRB constructs with small tails (compare Figure 3C and E). These results suggest that the long cytosolic C-terminal tail is important for both efficient insertion of strong hydrophobic C-TMD and flipping of less hydrophobic C-TMD into the ER lumen.

We next wanted to determine the minimum C-terminal cytosolic tail length required for either insertion of strong hydrophobic C-TMDs or flipping of insufficient hydrophobic C-TMDs. To this end, we prepared WRB (K164D) constructs with varying C-terminal length and tested them for insertion or flipping by metabolic labeling and immunoprecipitation. We found ~100 residues at the C-terminus rendered efficient flipping of C-TMD with insufficient hydrophobicity (Figure 3F). In contrast, about 60 residues at the C-terminus were sufficient to prevent flipping of C-TMDs with strong hydrophobicity (Figure 3F). This result is consistent with the model that the presence of the ribosome at the translocon is crucial for efficient insertion of C-TMDs with strong hydrophobicity, presumably by trigging the opening of the lateral gate of the translocon (Figure 3G). Collectively, these findings suggest that hydrophobicity alone is not sufficient for insertion of a TMD and that the C-terminal tail length also influences the insertion of a TMD into the ER membrane.

### Insufficient hydrophobic C-TMDs with short tails enable slow flipping into the ER lumen

Since the flipped form of WRB is routed for degradation, we hypothesized that the flipping of C-TMD of WRB into the ER lumen may be slow to provide a time window for assembly with the partner protein CAML. We, therefore, monitored the dynamics of C-TMD flipping by performing pulse-chase experiments. The cells expressing WRB constructs with short C-terminal tails were briefly labeled and chased for three hours. Also, we treated cells with a p97 ATPase inhibitor during the chase period to prevent retrotranslocation and degradation of WRB, thus allowing us to quantify the flipped C-TMD without losing the signal from degradation. More than 20% of C-TMD of wild type WRB containing a positively charged residue flipped into the ER lumen even during labeling as shown by glycosylation, and that the remaining ~25% flipped post-translationally during 3 h of chase period (Figure 4A). The C-TMD containing a negative charge residue (K164D) also yielded a similar result (Figure 4B). In sharp contrast, the flipping of C-TMD of WRB (K164L) with sufficient hydrophobicity occurred only during labeling, but it did not flip post-translationally into the ER lumen during the chase period (Figure 4C). We then investigated whether the length of the C-tail influences the flipping dynamics of C-TMD into the ER lumen. In contrast to the short C-tail, about 40% of C-TMD of WRB-Venus flipped into the ER lumen during labeling, and it did not post-translationally flip into the ER lumen during the chase period (Figure 4D). Since a negative charge in a TMD significantly increases its free energy value than a positive charge residue (Figure 3A), more than 60% of the C-TMD of WRB (K164D)-Venus flipped into the ER lumen even during labeling. Interestingly, a small proportion of WRB (K164D)-Venus was post-translationally flipped into the ER lumen (Figure 4E). Similar to the short C-tail, the C-TMD of WRB (K164L)-Venus with sufficient hydrophobicity flipped only during labeling but did not flip post-translationally into the ER lumen during chase period (Figure 4F). Taken together our results suggest that both the hydrophobicity of C-TMD and the cytosolic tail length contribute to the slow flipping of C-TMD into the ER lumen.

**Figure 4.**
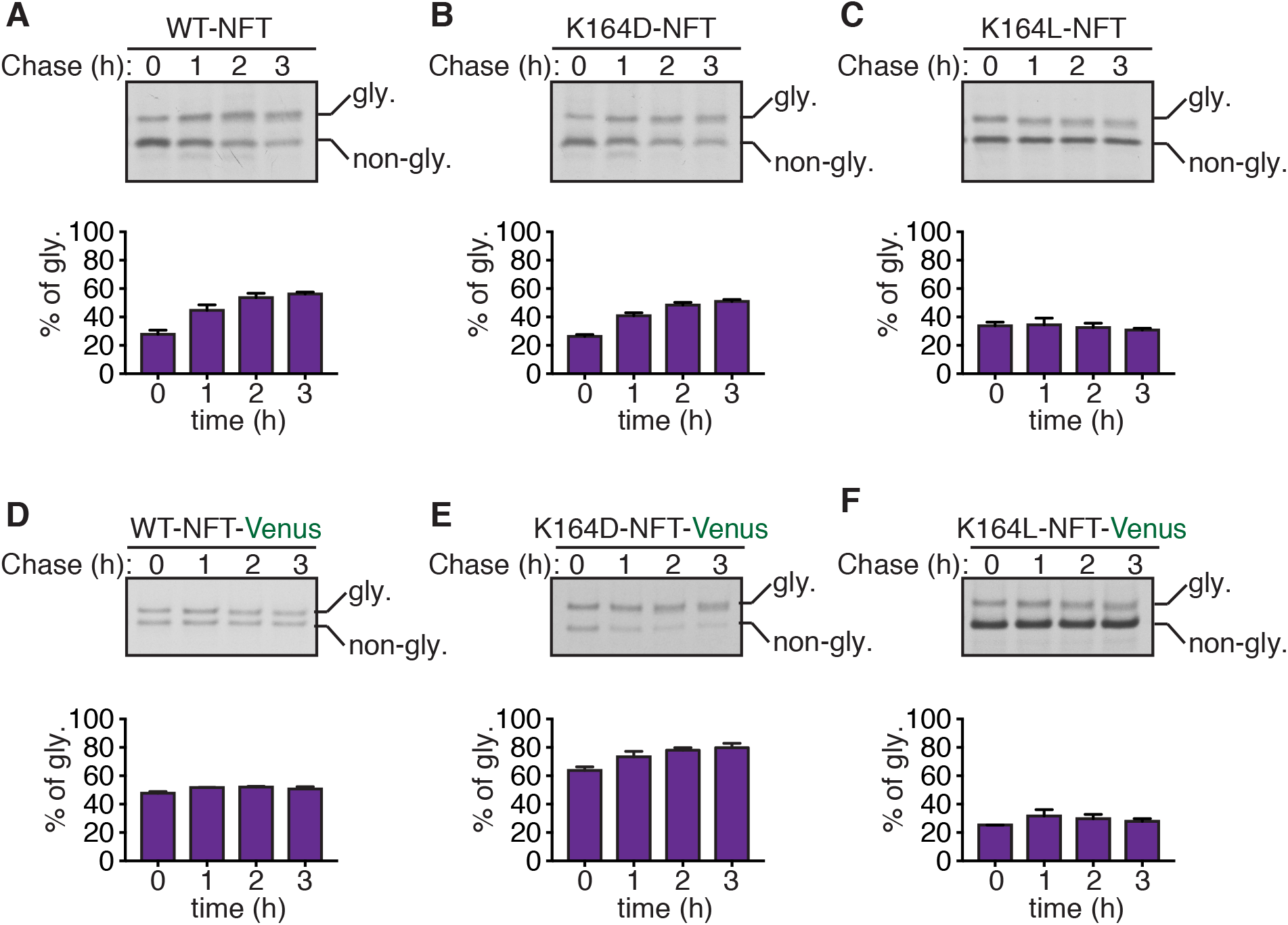
Insufficiently hydrophobic C-TMDs with short tails slowly flip into the ER lumen. (A) HEK293 cells were transfected with WRB-HA-NFT and metabolically labeled for 15 min and chased in the presence of a p97 ATPase inhibitor for indicated time points. Cell lysates were immunoprecipitated with anti-HA antibody beads and analyzed by autoradiography. The percentage of the flipped form quantified from three independent experiments and their standard deviations are depicted in a bar graph. (B)-(F) The indicated constructs were transfected and analyzed as in (A).

### The Sec61 translocon complex serves as a holdase for insufficiently hydrophobic C-TMDs to allow efficient assembly with partner TMDs

We hypothesized that the slow flipping of C-TMD of WRB may provide a time window to facilitate efficient assembly with its partner TMD from CAML (Figure 5A). Conversely, the C-TMD that either quickly inserts into membrane or flips into the ER lumen may not have sufficient time to efficiently assemble with its partner TMD from CAML. To test this, we performed CAML pulldown with WRB constructs that vary in their hydrophobicity of C-TMDs. We noticed that coexpression of CAML with WRB reduced the flipped form of WRB compared to when WRB was expressed alone, suggesting that CAML can prevent the flipping of C-TMD into the ER lumen (Figure 5B, input). Indeed, CAML selectively co-immunoprecipitated with the non-glycosylated inserted form of WRB (Figure 5B). Strikingly, the assembly of CAML with WRB (K164L), the C-TMD of which quickly flipped (Figure 4C), was significantly lower than wild type WRB (Figure 5B). Surprisingly, replacing lysine with a negatively charged residue in the C-TMD of WRB also assembled with CAML similar to the wild type (Figure 5B). To exclude the possibility that the replacement of lysine to leucine in the C-TMD of WRB interfered its assembly with CAML, we tested the interaction between purified CAML and WRB protein in vitro. In contrast to the observation from cells, CAML efficiently interacted with WRB (K164L) compared to the wild type (Figure 5C). This result emphasizes that the lysine residue in the C-TMD is not directly involved in the assembly with CAML, but rather it facilitates the slow flipping of the TMD into the ER lumen. We then investigated the effect of the C-terminal length of WRB on assembly with CAML. We hypothesized that WRB-Venus may not assemble efficiently with CAML since the larger C-terminal Venus tag facilitated a faster flipping of the C-TMD into the ER lumen (Figure 4E-F). To test this, we compared CAML assembly with either WRB constructs with short tails or WRB constructs with large Venus tag. In support of our hypothesis, CAML efficiently assembled with WRB containing short C-tails, but inefficiently assembled with WRB containing large Venus tag (Figure S5). We next asked whether the C-terminal length-dependent flipping observed in Figure 3F would correlate with the assembly efficiency with CAML. To address this, we performed interaction studies between CAML and WRB constructs harboring varied length of C-terminal tails. This result further supported our view that slow flipping WRB constructs that contain less than 84 residues at their C-terminus assembled efficiently with CAML (Figure 5D). In sharp contrast, faster flipping WRB constructs that contain more than 100 residues at their C-terminus very poorly assembled with CAML. This conclusion is further corroborated by the observation that CAML levels were also stabilized upon co-expression of WRB constructs containing less than 84 residues at their C-terminus (Figure 5D).

**Figure 5.**
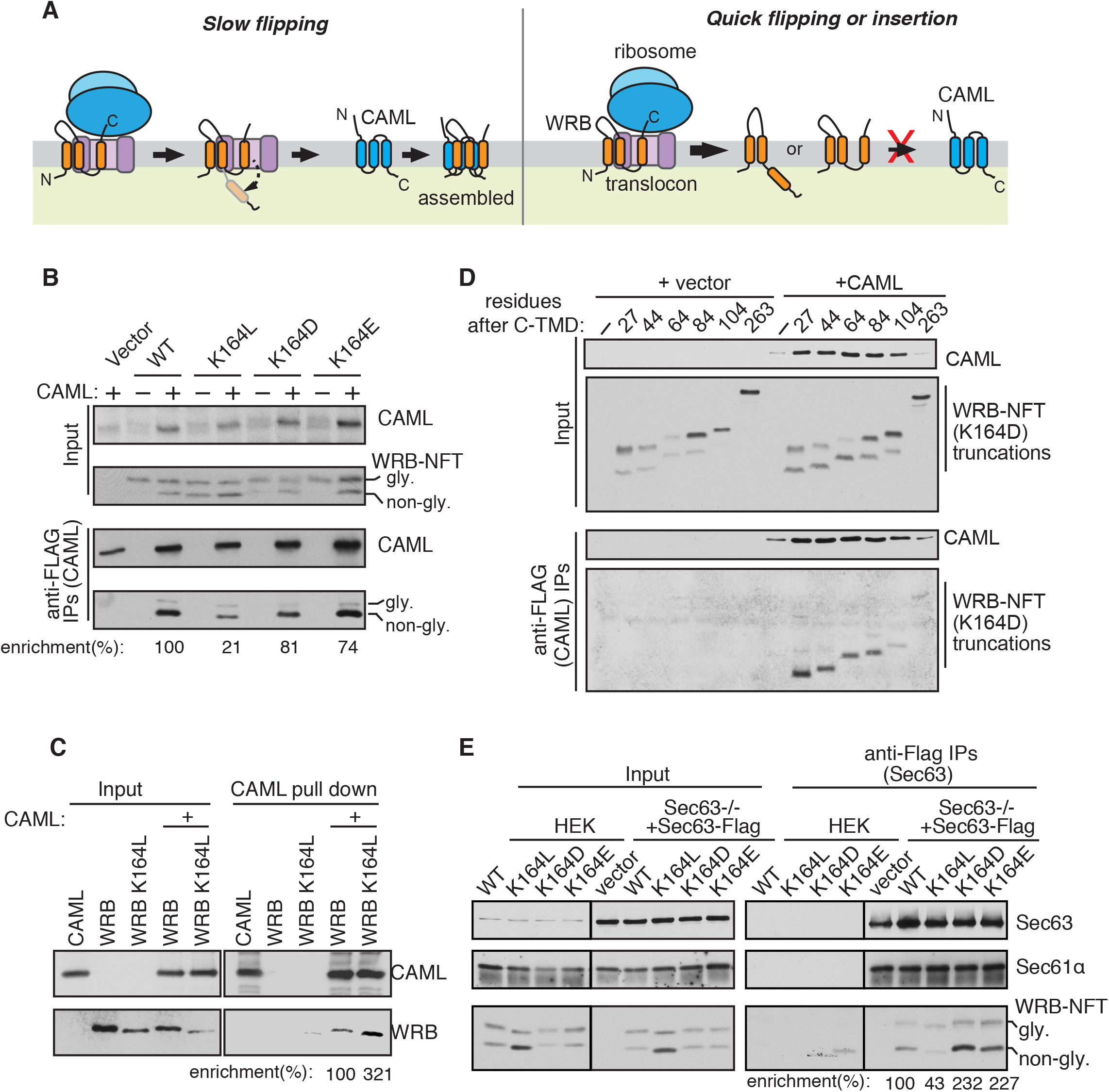
The C-TMD flipping rate determines the assembly efficiency with its partner TMD. (A) Left panel: A hypothetic model showing the slow flipping of insufficiently hydrophobic C-TMD provides a time window for the efficient assembly with the partner TMD. Right panel: The C-TMD that either quickly inserts into the membrane or flips into the ER lumen poorly assembles with the partner TMD. (B) The indicated WRB-HA-NFT constructs were co-transfected with CAML-FLAG into HEK293 cells and immunoprecipitated with anti-FLAG beads. The resulting samples were analyzed by immunoblotting with anti-FLAG antibody (CAML) and anti-HA antibody (WRB). The band intensity was quantified by Image J, and the ratio of IPs to inputs was calculated. The IP enrichment of wild type WRB was taken as 100%. (C) WRB-FLAG or WRB (K164L)-FLAG transiently transfected and purified from 293T cells using anti-FLAG beads. CAML-HA was expressed in 293T cells and immunoprecipitated with anti-HA beads. The purified WRB-FLAG or WRB (K164L)-FLAG was mixed with CAML-HA bound to anti-HA beads. The beads were washed and the bound proteins were eluted with SDS sample buffer and analyzed by immunoblotting with anti-FLAG (WRB) and anti-HA (CAML) antibodies. The percentage of enrichment was calculated as in (B). (D) The indicated WRB-NFT-Venus truncations were co-transfected with CAML-FLAG into HEK293 cells and analyzed as in (B). (E) The indicated WRB-HA-NFT variants were transfected into Sec63−/− HEK293 cells stably expressing Sec63-FLAG and immunoprecipitated with anti-FLAG beads. The resulting samples were analyzed by immunoblotting with Sec63 antibodies, Sec61α antibodies, and HA antibodies (WRB-HA-NFT variants). The percentage of enrichment was quantified as in (B).

These findings thus far suggest that insufficient hydrophobic C-TMDs with short tails are transiently retained by a membrane holdase and are flipped into the lumen when they failed assembly with partner TMDs. We hypothesized that the Sec61 translocon may be a post-translational TMD holdase since it is responsible for recognizing the C-TMD right after its release from the ribosome. To selectively enrich the post-translationally retained C-TMD in the Sec61 translocon, we used the translocon associated membrane protein Sec63 as a handle since it occludes the ribosome binding to the translocon (Itskanov and Park, 2019; Wu et al., 2019). Although Sec63 is known to facilitate post-translational translocation of proteins into the ER lumen (Deshaies et al., 1991; Matlack et al., 1999; Panzner et al., 1995; Wu et al., 2019), it is dispensable for the C-TMD of WRB flipping into the ER lumen (Figure S6). As expected, Sec63 formed a complex with the Sec61 translocon as it was efficiently co-immunoprecipitated with the translocon (Figure 5E). In support of our hypothesis, C-TMDs of WRB constructs with insufficient hydrophobicity were enriched with the Sec63/Sec61 translocon complex. In contrast, the C-TMD (K164L) with sufficient hydrophobicity was weakly associated with the translocon even though it expressed higher than other constructs (Figure 5E), thus explaining its poor assembly with CAML. Taken together our results support a model in which insufficient hydrophobicity C-TMDs with short tails are retained by the translocon, thus providing a sufficient time window for the assembly with partner TMDs.

### The C-terminal length determines localization and degradation of orphaned membrane proteins

We next investigated the mechanism by which the flipped C-TMD is recognized by the ERAD pathway. We tested the role of the conserved E3 ligase Hrd1 in recognition of unassembled WRB constructs with shorter C-terminal tails. To our surprise, the pulse-chase experiment revealed that WRB degradation was less dependent on Hrd1 E3 ligase as WRB was similarly turned over both in control HEK293 cells and Hrd1−/− HEK293 cells created by CRISPR/Cas9 (Figure 6A, B). To directly monitor the turnover of the flipped C-TMD, we analyzed our glycosylation reporter construct of WRB by the pulse-chase experiment. The degradation of the flipped form of WRB was also less dependent on the activity of Hrd1 E3 ligase as it was turned over quickly both in wild type and Hrd1−/− cells (Figure 6C). We also obtained a similar result when we replaced the lysine residue in the C-TMD with aspartic acid (Figure S7A). We further tested whether Hrd1 independent degradation of the flipped TMD with a short tail applies to other unassembled membrane proteins. Indeed, the degradation of both flipped C-TMDs of TRAP*γ* and PMP22 did not require the activity of Hrd1 E3 ligase (Figure S7B-E).

**Figure 6.**
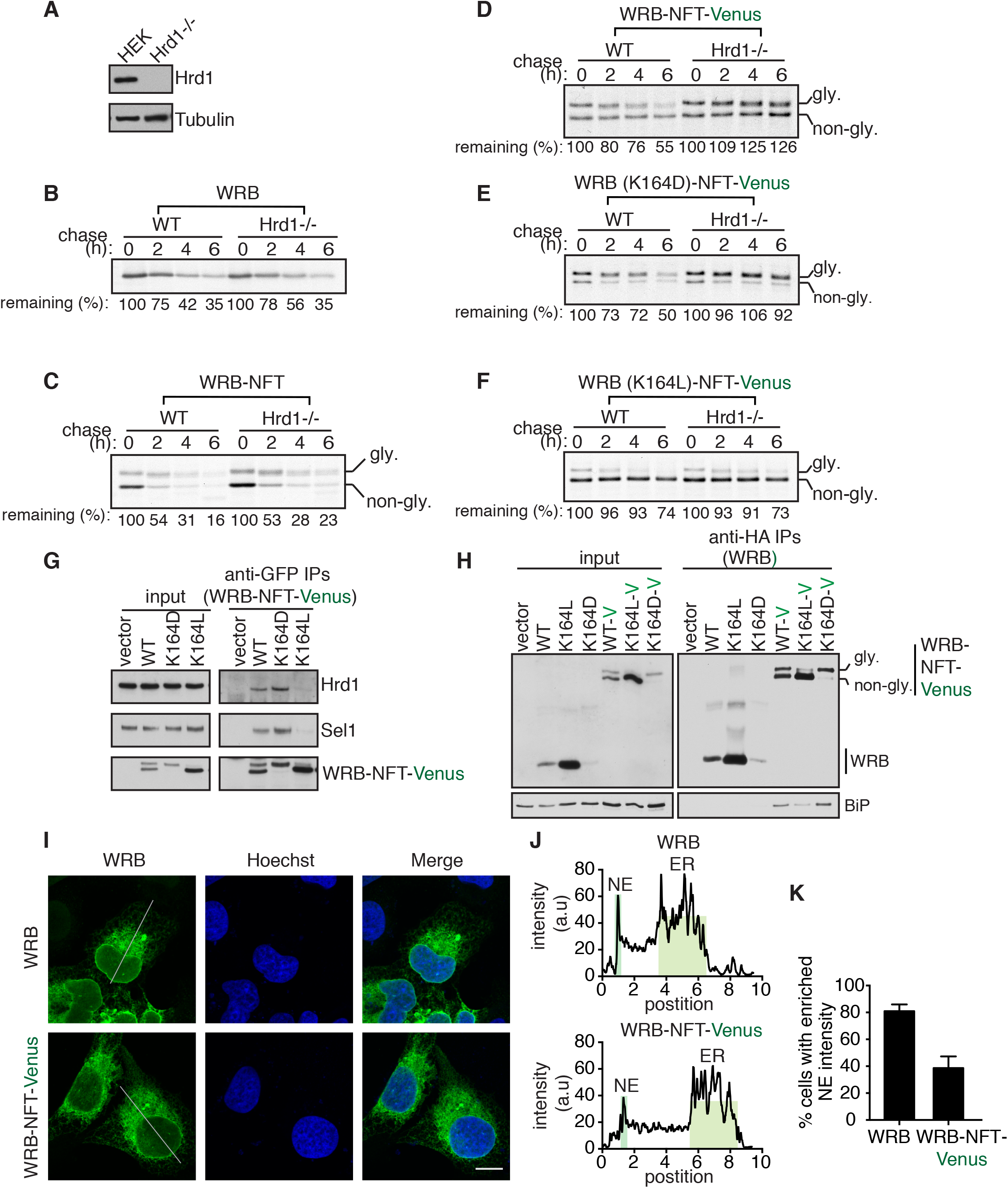
The C-terminal length determines the localization and degradation of unassembled membrane proteins. (A) The lysate of either HEK293 or Hrd1−/− HEK293 cells was immunoblotted for the indicated antigens. (B) HEK293 or Hrd1−/− cells expressing WRB-HA were metabolically labeled and chased for the indicated time points. The lysates were analyzed by autoradiography after immunoprecipitation with anti-HA beads. The protein level at 0-hour time point was taken as 100%, and the percentage of the remaining protein was calculated with respect to 0 hour. (C)-(F) The indicated cell lines were transfected with the indicated WRB variants and analyzed as in (B). (G) The indicated WRB-NFT-Venus variants were transfected into HEK293 cells and immunoprecipitated with anti-GFP antibodies and immunoblotted for the indicated antigens. (H) The indicated WRB-HA and WRB-HA-NFT-Venus variants were transfected into HEK293 cells and immunoprecipitated with anti-HA beads, and analyzed by immunoblotting with anti-BiP antibody, and anti-HA antibodies for variants of WRB. (I) U2OS cells were transfected with WRB-HA or WRB-NFT-Venus. After 24 hours of transfection, the cells were processed for immunostaining with anti-HA or anti-GFP antibodies Representative images of three independent experiments are shown. Scale bar: 10μm. (J) The line intensity scans from images (I) were plotted. Note that WRB-HA has a higher intensity on the nuclear envelope compared to WRB-NFT-Venus. (K) The number of cells enriched with NE intensity for WRB-HA and WRB-NFT-Venus were quantified by Image J. The quantification from three independent experiments and their standard deviations are depicted in a bar graph.

To investigate whether the C-terminal length of the flipped TMD determines its recognition by Hrd1 E3 ligase, we analyzed the turnover of WRB-Venus. Remarkably, the degradation of the flipped C-TMD with the large Venus tag strictly relied on Hrd1 E3 ligase since the flipped form of WRB-Venus was completely stabilized in Hrd1−/− cells compared to wild type cells (Figure 6D). The degradation of the C-TMD of WRB-Venus containing aspartic acid instead of lysine was also dependent on Hrd1 activity (Figure 6E), arguing that Hrd1 E3 ligase does not seem to recognize a specific charge residue in the flipped TMD, but rather recognizes the flipped TMD with the Venus tag. Replacing lysine to leucine increased the hydrophobicity of C-TMD of WRB-Venus, which in turn significantly reduced the flipping C-TMD into the ER lumen (Figure 6F). This construct was relatively stable in both wild type and Hrd1−/− cells (Figure 6F). To ascertain our findings, we repeated pulse chase experiments of WRB constructs using Sel1−/− cells, which is a subunit of the Hrd1 E3 ligase complex. Consistent with our results from Hrd1−/− cells, we found that the turnover of the flipped C-TMD with the Venus tag was dependent on Sel1, whereas the degradation of the flipped C-TMD with a short tail was independent of the Sel1 activity (Figure S8).

Since WRB constructs with a large C-terminal Venus tag are degraded in a Hrd1 E3 ligase dependent manner, we assessed whether it can associate with Hrd1 E3 ligase by the coimmunoprecipitation assay. Consistent with our protein turn over data, we found that Hrd1 selectively interacted with C-TMD containing charged residues (WRB or WRB (K164D)) but not with sufficiently hydrophobic C-TMD (K164L), suggesting that the efficient flipping of C-TMD-Venus into the ER lumen is required for the binding with Hrd1 (Figure 6G). We next asked how Hrd1 E3 ligase selectively recognizes the long C-tail of the flipped TMD, but not the C-TMD with short tails. Since the luminal chaperone BiP is known to capture luminal misfolded proteins and deliver them to the Hrd1 E3 ligase complex, we hypothesized that BiP may selectively bind to the flipped TMD with a large C-terminal domain. To test this, we co-immunoprecipitated WRB constructs with short and long C-tails and probed for their association with BiP. In support of our hypothesis, BiP only interacted with WRB containing the large C-terminal Venus tag but not with WRB containing shorter tails. This interaction likely occurs in the ER lumen since WRB (K164L), which does not flip efficiently, interacted weakly with BiP compared to the wild type and K164D-Venus (Figure 6H). Finally, we investigated the role of C-terminal length in the localization of unassembled membrane proteins. Interestingly, confocal imaging of WRB or WRB-Venus revealed that WRB with a short C-tail decorated the nuclear envelope in addition to the ER localization. By contrast, most WRB-Venus was localized in the ER with a small proportion localized in the nuclear envelope (Figure 6I-K). These results implicate that unassembled C-TMDs with short tails can freely diffuse into the nuclear envelope since their inability to bind with BiP. In contrast, BiP efficiently captures C-TMDs with large domains and delivers them to the Hrd1 E3 ligase complex for retrotranslocation into the cytosol for the proteasomal degradation.

## DISCUSSION

A large proportion of membrane proteins exist in multi-membrane protein complexes and they serve fundamental roles in the cell physiology (Juszkiewicz and Hegde, 2018). However, it is poorly understood how these newly synthesized membrane proteins find each other in the crowed and extensive network of the ER membrane, which includes many competing factors such as chaperone-linked quality control components (Brodsky, 2012; Wu et al., 2018). In this study, we uncover multiple new roles for the C-terminal tails of membrane proteins in insertion, assembly and quality control of membrane proteins (Figure 7).

**Figure 7.**
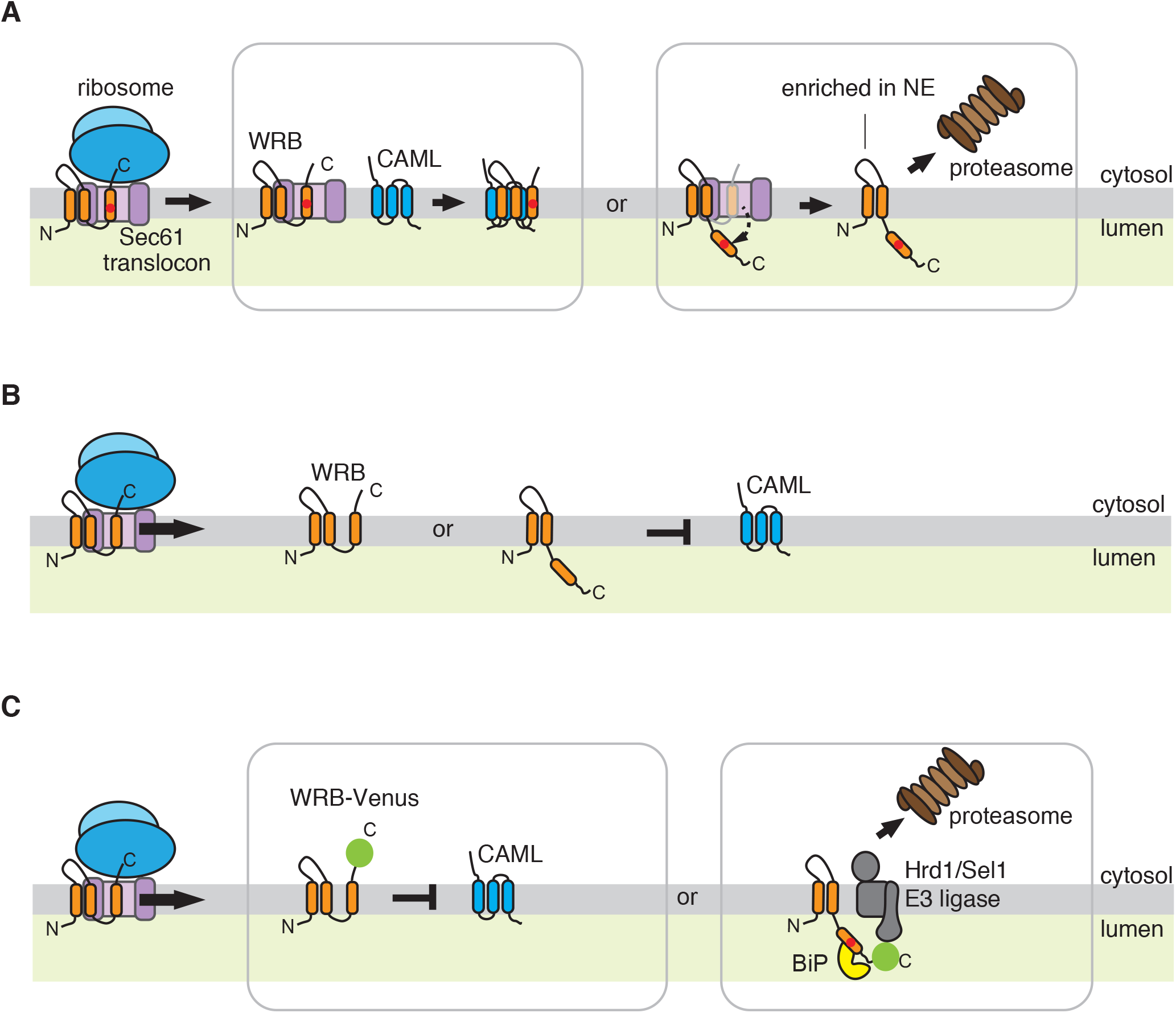
Models showing the roles of the C-terminal tail in insertion, assembly and degradation of membrane proteins. (A) Insufficiently hydrophobic C-TMDs with short cytosolic tails are post-translationally retained by the Sec61 translocon. The Sec61 mediated retention of C-TMDs provide a time window to assemble with partner TMDs. If the C-TMD is unassembled, it slowly flips through the translocon and can move to the nuclear envelope. (B) Sufficiently hydrophobic C-TMDs with short cytosolic tails disobey the biological hydrophobicity scale. They are quickly either inserted into the membrane or flipped into the ER lumen, resulting in a much shorter residence time in the translocon for the assembly with partner TMDs. (C) C-TMDs with long cytosolic tails follows the biological hydrophobicity scale. Accordingly, C-TMDs with insufficient hydrophobicity quickly flips into the ER lumen, whereas C-TMDs with strong hydrophobicity quickly insert into the ER membrane. Thus, the C-TMD does not reside long enough in the translocon for efficient assembly with the partner TMD. The flipped C-TMD with long cytosolic tail is captured by the luminal chaperone BiP. The BiP bound C-TMD is then subsequently recognized by the Hrd1/Sel1 E3 ligase complex for retrotranslocation and degradation by the proteasome.

To reveal the biochemical features that are necessary for both membrane protein assembly and degradation, we chose to investigate the assembly of the tail-anchored membrane protein complex comprising WRB and CAML proteins because of their smaller sizes, which can be biochemically manipulated. Also, their assembly is mediated mainly through their transmembrane domains (Vilardi et al., 2014). While WRB contains three TMDs with N-terminus in the lumen and C-terminus in the cytosol, CAML has three TMDs with N-terminus in the cytosol and C-terminus in the lumen (Figure 7). Both proteins are robustly degraded when they failed to assemble in the ER membrane. However, coexpression of both membrane proteins increases their stability by shielding from recognition quality control factors for ubiquitination and retrotranslocation into the cytosol. By performing a detailed domain mapping and mutagenesis studies, we identify that a single lysine residue in the C-TMD of WRB is responsible for flipping the TMD into the ER lumen in the absence of partner protein. A recent study suggests that the second and C-TMD of CAML also flipped into the ER lumen in the absence of WRB (Carvalho et al., 2019). Although it remains to be determined how WRB fixes the orientation of CAML, this study also suggest that the C-TMD flipping is a prevalent mechanism for unassembled polytopic membrane proteins. This notion is further supported by our data that either the C-TMD of TRAPγ or Sec61α flip into the ER lumen when it fails to assemble with the partner protein. Our data derived from introducing a patient mutation in PMP22 suggested that C-TMDs of polytopic membrane proteins that do not assemble with their neighboring TMDs can also flip into the ER lumen.

Although the TMD flipping into the ER lumen has been reported by previous studies(Buck and Skach, 2005; Coelho et al., 2019; Feige and Hendershot, 2013; Kim and Skach, 2012; Rabeh et al., 2012; Roushar et al., 2019), the biochemical features responsible for flipping remain unclear. We find that the flipping of C-TMD is not just due to less hydrophobic TMD, but rather its C-terminal tail length determines the flipping of the TMD. Even a strong hydrophobic C-TMD does not efficiently insert into the membrane and flips into the ER lumen if the C-terminus contains less than 60 amino acids. This is likely caused by the termination of translation before the C-TMD releases from the ribosome exit tunnel, which can accommodate ~40 residues (Voss et al., 2006). Hence, the C-TMD is post-translationally recognized by the Sec61 translocon. However, the post-translational insertion of C-TMDs with short tails are inefficient because the lateral gate of the translocon is likely closed in the absence of the ribosome (Voorhees et al., 2014; Voorhees and Hegde, 2016a, b; Wu et al., 2019).

The C-terminal tail length plays a unique function in the case of C-TMDs with insufficient hydrophobicity. As expected, these hydrophilic C-TMDs flips into the ER lumen, but their flipping was more efficient when their C-terminal was extended longer than 100 residues. Our pulse-chase experiments revealed that insufficiently hydrophobic C-TMDs with short tails exhibit a slow flipping rate. This result is consistent with the previous study analyzing a single spanning membrane protein TCRα, which continuously post-translationally enters into the ER lumen (Feige and Hendershot, 2013). These data suggested that the slow flipping of less hydrophobic C-TMDs into the ER lumen may provide a time window for the assembly with the partner protein. Indeed, C-TMDs of WRB that slowly flips into the ER lumen efficiently assemble with CAML. Previous studies have shown that a charged residue in a TMD of membrane protein is often required for making an ionic bond with an opposite charge in the partner TMD of membrane protein (Cosson et al., 1991; Feige and Hendershot, 2013; Roushar et al., 2019). Our data suggest that charged residues in TMDs may also be evolved to interact with membrane protein holdases, thus allowing efficient assembly with partner TMDs.

A large proportion of newly synthesized insufficiently hydrophobic C-TMDs with short tails is initially localized in the membrane and slowly flips into the ER lumen. This data suggested that the Sec61 translocon may post-translationally hold these TMDs. Indeed, we can demonstrate that insufficiently hydrophobic C-TMDs can interact with the Sec61 translocon complex better than the C-TMD with sufficient hydrophobicity. The interaction between insufficiently hydrophobic C-TMDs of WRB and the Sec61 translocon correlates with the assembly efficiency with CAML. It remains to be understood how CAML pulls the C-TMD of WRB from the Sec61 translocon. It is possible that the first TMD of WRB, which is likely outside the translocon, may recruit CAML to the Sec61 translocon. Interestingly, the endogenous level of CAML is estimated to be five times higher than WRB (Colombo et al., 2016), thus suggesting that CAML has a higher chance to release WRB from the translocon complex.

We propose that our findings of C-terminal length influencing insertion, flipping and assembly of membrane proteins are important to understand the biogenesis of glycosylphosphatidylinositol (GPI) anchored proteins (Kinoshita and Fujita, 2016). The GPI anchor sequences are typically localized at the C-terminus of GPI anchored proteins. These GPI anchored sequences are moderately hydrophobic with no or very short C-terminal tails. Similar to C-TMD of WRB, the GPI anchor sequences are likely post-translationally recognized and slowly flipped into the ER lumen. This view is supported by the previous studies that GPI anchored sequences cannot be inserted into the membrane even when extending their C-terminus with additional cytosolic sequences (Dalley and Bulleid, 2003). It is tempting to suggest that GPI anchored sequences also post-translationally retained by the Sec61 translocon, thus providing a kinetic time window for the GPI anchor addition to the C-terminal end of a GPI anchored protein by the GPI transamidase complex. However, this hypothesis needs to be tested in future studies.

The unassembled C-TMDs of membrane proteins are slowly flipped into the ER lumen and are recognized by quality control factors for retrotranslocation and degradation by cytosolic proteasomes. The flipped TMDs with short C-terminal tails are not efficiently recognized by the conserved the Hrd1/Sel1 E3 ligase complex. In sharp contrast, the flipped TMDs with long C-terminal tail strictly rely on recognition by Hrd1 in the ER membrane. Our data suggest that C-terminal length-dependent recognition can be attributed to BiP, which selectively binds to the flipped TMDs with long C-terminal tail and delivers them to the Hrd1 complex. This observation is consistent with the previous studies where the completely translocated TCRα is captured by BiP and delivered it to Hrd1(Feige and Hendershot, 2013). It remains to be addressed why BiP cannot bind to the flipped TMD with a short tail. A possibility is that the flipped TMD with a short tail is very proximal to the ER membrane, thus recognition by BiP is challenging. If BiP is unable to bind to the flipped TMD, it can freely move to the nuclear envelope and presumably undergoes degradation by the inner nuclear membrane quality control pathway. Indeed, it has been shown that many unassembled ER membrane proteins are often mislocalized to the inner nuclear membrane and are recognized by the Asi E3 ligase complex for degradation in yeast (Foresti et al., 2014; Khmelinskii et al., 2014). However, the mammalian homolog of Asi E3 ligase complex remains to be identified. Polar amino acid mutations in TMDs of membrane proteins are frequently associated with human diseases such as retinitis pigmentosa, cystic fibrosis, and Charcot-Marie-Tooth disease (Partridge et al., 2004). It is tempting to suggest that these membrane proteins with insufficiently TMDs may be transiently trapped within the translocon, thus causing ER stress and apoptosis, which can alter disease phenotypes.

## STAR METHODS

Detailed methods are provided in the online version of this paper and include the following:

- KEY RESOURCES TABLE
- CONTACT FOR REAGENT AND RESOURCE SHARING
- EXPERIMENTAL MODEL AND SUBJECT DETAILS
  - Cell lines
- METHOD DETAILS
  - Constructs
- QUANTIFICATION OF STATISTICAL ANALYSIS

## SUPPLEMENTAL INFORMATION

Supplemental information includes nine figures and can be accessed with this article online at …

## ACKNOWLEDGMENTS

We thank Dr. Peter Cresswell and Dr. Jeffrey Grotzke for Hrd1 gRNA and Hrd1−/− HEK293 cells. We are grateful to Dr. Ramanujan Hegde for Sec61alpha and Sec63 antibodies. We want to thank Dr. Ling Qi for mouse Sel1 plasmid. We thank Dr. Zai-Rong Zhang for comments on the manuscript. We thank Jacob Culver and Dr. Xia Li for discussions and comments on the manuscript. This work was supported by NIH 1R01GM117386-01A1 and 1R21AG056800-01A1.

## AUTHOR CONTRIBUTIONS

S.S. performed the majority of the experiments with help from M.M. The project was supervised by M.M. The manuscript was written by S.S. and M.M.

## DECLARATION OF INTERESTS

The authors declare no competing interests.

## METHOD DETAILS

### DNA constructs

All the constructs were created using the pcDNA/FRT/TO vector (Invitrogen). WRB-FLAG, WRB-HA, WRB-HA-NFT, WRB-FLAG-AP, CAML-FLAG, CAML-HA, CAML-HA-NFT, PMP22-HA, PMP22-HA-NFT, TRAPγ-HA, TRAPγ-HA-NFT, Sec61α-HA, and Sec61α-HA-NFT were generated using standard molecular biology methods. pcDNA/FRT/TO containing the fusion of WRB and CAML was constructed by including a 15 residues linker (ASGAGGSEGGTSGAT) between these genes using the overlap PCR method. pcDNA/FRT/TO containing WRB-TMD1-swap-FLAG was constructed by replacing WRB TMD1 (amino acids 9-29) with yeast Ost4pTMD (amino acids 10-28). WRB-TMD2-swap-FLAG was cloned by replacing WRB TMD2 (amino acids 100-120) with human Sec61β TMD (amino acids 71-91). WRB-TMD3-swap-FLAG was generated by replacing WRB TMD3 (amino acids 149-169) with yeast Ost4p TMD (amino acids 10-28). WRB-HA-NFT-Venus was created by overlap PCR with standard molecular biology methods. While the Pfu polymerase (Agilent Technologies) was used for the site directed mutagenesis, the Phusion polymerase (NEB) was used for other PCR reactions. pCDNA3 HA-Ubiquitin plasmid was described previously (Zhang et al., 2013). The coding sequences of all constructs were verified by sequencing (Yale Keck DNA Sequencing) to preclude any sequence error.

### Cell culture and the generation of CRISPR/Case9 knock out cells

293T and HEK 293-Flp-In T-Rex cells (kindly provided by Dr. Ramanujan Hegde, MRC, UK) were cultured in high Glucose Dulbecco’s Modified Eagle’s Medium (DMEM) with 10% fetal bovine serum (FBS) and 100 U/mL penicillin and 100 μg/mL streptomycin at 5% CO_2_. Hrd1−/−, Sel1−/−, or Sec63−/− HEK 293-Flp-In T-Rex cell lines were generated using the CRISPR/Cas9 system as previously described (Ran et al., 2013, Mali et al., 2013).. HEK293-Flp-In T-Rex cells were transfected with pSpCas9(BB)-2A-Puro and gRNA expression plasmid of Hrd1(AGAGTGCAACAAAGCGG), Sel1(ACTGCAGGCAGAGTAGTTGC), or Sec63(GCCAGAGGTAGTATGTCGC). Cells were grown for 24 h and treated with 2.5μg/mL puromycin for 72h to select the successfully transfected clones. Single-cell clones were isolated by plating at 0.5 cells/well in 96 well plates. Hrd1, Sel1, and Sec63 knock outs were confirmed by immunoblotting. All the cell lines used in this study were not tested for mycoplasma, but many cell lines were used in immunofluorescence assays with Hoechst staining that should reveal presence of mycoplasma. Cells were assumed to be authenticated by their respective suppliers and were not further confirmed in this study. However, knock-out lines were routinely validated by immunoblotting.

To establish Sec63−/− HEK293 cells stably expressing Sec63-FLAG, 1.6μg of pOG44 vector (Invitrogen) and 0.4μg of pcDNA/FRT/TO-Sec63-FLAG were transfected into Sec63−/− cells in a well in the 6 well plate using Lipofectamine 2000 (Invitrogen). After 24h of transfection, cells were transferred to 10 cm dishes and selected with 150 μg/ml hygromycin (Invitrogen) and 10μg/ml blasticidin (InvivoGen, San Diego, CA). The selection medium was replaced every three days until colonies appeared. The colonies were picked and the protein expression was evaluated by immunoblotting. Transfections in HEK 293-Flp-In T-Rex cells were performed using Lipofectamine 2000 (Invitrogen) according to the manufacture’s protocol but with either half or quarter amount of the recommended DNA and Lipofectamine 2000.

### Cycloheximide chase

Wild type HEK 293-Flp-In T-Rex or Hrd1−/− cells (0.15 × 10^6^/well) were plated on polylysine pre-coated (0.15mg/mL) 24-well plates and transiently transfected with 400ng or 800ng (for co-expression experiments, 400ng of each plasmid) of the indicated construct in figures. Typically, expression of indicated proteins were induced with doxycycline (200ng/mL) for 24 hours prior to treatment with cycloheximide (150μg/ml). The treated cells were directly collected with 100ul of 2x SDS sample buffer at indicated time points. In some cases, cells were treated both cycloheximide and MG132 (20μM). Samples were then analyzed by immunobloting with indicated antibodies in the Figure legends.

### Subcellular fractionation

To assay the retrotranslocation of WRB and CAML from the ER to the cytosol, HEK 293-Flp-In T-Rex cells (0.6 × 10^6^ /well) were transiently transfected with 2μg indicated plasmids in the figures and induced with 200ng/mL doxycycline. After 24 hours of transfection, cells were treated with either 40μM MG132 alone or 10μM P97 inhibitor (NMS-873, Millipore sigma) (Magnaghi et al., 2013) plus 40μM MG132 or equal amount of DMSO for 2 hours. Cells were then washed twice with KHM buffer (20mM Hepes pH7.4, 110mM KAC, 2mM MgAC), and incubated with 800μL KHM contained 0.005% digitonin for 5 min on ice to permeablize the cell membrane. The cytosol fraction was collected and cleared by centrifugation at 12000rpm for 2 min. The membrane fraction that was left on the plate was washed once with KHM buffer and harvested in 800uL RIPA buffer followed by centrifugation at 20,000g for 15 min. 50μL of samples were taken from both the cytosol and the membrane fractions and mixed with 25μL 5x SDS sample buffer. The samples were analyzed by immunoblotting with anti-FLAG antibody.

### Glycosylation assay and Endo H treatment

HEK 293-Flp-In T-Rex cells (0.15 × 10^6^ /well) were plated on poly-lysine (0.15mg/mL) coated 24-well plates and transiently transfected with 400ng indicated plasmids. Protein expression was induced with 200ng/mL doxycycline. After 24 hours transfection and induction, cells were collected with 60μL SDS Buffer (1% SDS, 0.1M Tris pH 8.0). The lysates were diluted with 60μL of 2%BME, 0.1M Tris (pH 6.8) and boiled at 100°C for 5 min. The samples were then divided into two aliquots (60μL for each), and were treated with or without 2μL (1000 units) of Endo H (NEB) in a 100μL reaction volume including 1x G5 buffer (NEB) at 37°C for 4 hours. The reaction was terminated by adding 50μL 5X SDS sample buffer and boiled for 5 min before analysing by immunoblotting with anti-HA antibody.

### Metabolic Labeling and pulse chase assay

HEK 293-Flp-In T-Rex or Hrd1−/− cells (0.6 × 10^6^/well) were plated on polylysine coated (0.15mg/mL) 6-well plates and transiently transfected with 2μg indicated plasmids. Expression of indicated proteins was induced with doxycycline (200ng/mL) for 24 hours prior to the chase. Cells were starved with non-Met/Cys medium including 10% dialyzed FBS and doxycycline for 30 min. For metabolic labeling assay in Figure 3, cells were labeled with 80μCi/mL Express^35^S protein labeling Mix for 15min. For pulse chase assays, cells were labeled with the ^35^S labeling mix for 30 min and chased in DMEM medium supplemented with 2mM Methionine and 2mM Cysteine for indicated time. For Figure 4, 10μM P97 inhibitor (NMS-873, Millipore Sigma) was added during starvation, labeling and chase periods. The labeled cells were collected and lysed in RIPA buffer including a protease inhibitor cocktail (Roche). The lysate was cleared by centrifugation at 20,000g for 15 min. The supernatant was mixed with either anti-HA magnetic beads or anti-GFP antibodies for 2 hours with rotation in the cold room. After incubation, protein-A agarose was added to the samples containing anti-GFP antibodies and incubated for 1 hour. The immunoprecipitates were washed three times with 1 ml of RIPA buffer, and proteins were eluted with 2X SDS sample buffer and run on 7.5% or 10% Tris-Tricine SDS-PAGE gels. The gels were dried and analyzed by autoradiography.

### *In vitro* translation and insertion

Transcripts encoding WRB-NFT-HA variants were obtained from *in vitro* transcription reactions as described previously (Mariappan et al., 2010). The transcripts were translated in the presence or absence of canine pancreas rough microsomes for 45 min at 32°C. The translated products (10ul) were denatured with 100ul of Tris-SDS buffer (0.1M Tris-HCl and 1% SDS) and diluted with the Triton buffer (50mM Tris-HCl pH 8.0, 150mM NaCl and 1% TritonX-100) before immunoprecipitating with anti-HA magnetic beads. The resulting samples were analyzed by SDS-PAGE and autoradiography.

### Immunoprecipitation

To examine the polyubiquitination of WRB or CAML, HEK 293-Flp-In T-Rex cells (0.6 × 10^6^ /well) were plated on poly-lysine (0.1mg/mL) coated 6-well plates and transiently transfected with 500ng pCDNA3/HA-Ubiquitin plasmid with 1.5ug of the indicated constructs in the Figure 1 After 24 hours transfection, cells were harvested in RIPA buffer containing a protease inhibitor cocktail (Roche) and 2mM N-ethylmaleimide. After centrifugation at 20,000g for 10 min, the lysates were incubated with rat anti-FLAG beads (Biolegend) for 1.5 hours. The beads were then washed with 1ml of RIPA buffer for 3 times. The washed beads were directly treated with 50μL 2x SDS sample buffer and boiled for 5 minutes followed by immunoblotting with anti-HA and anti-FLAG antibodies.

To probe the interaction between CAML and WRB variants in Figure 5, HEK 293-Flp-In T-Rex cells (0.6 × 10^6^ /well) were plated on poly-lysine coated 6-well plates and transiently transfected with 1μg pcDNA/FRT/TO-CAML-FLAG and 1μg pcDNA/FRT/TO-WRB-HA variants and induced the protein expression by treating cells with 200ng/mL doxycycline. After 24 hours transfection, cells were washed and harvested in PBS. The cell pellets were obtained by centrifuging for 2 min at 10,000g. The cell pellets were lysed with 200μL Buffer A (2% digitonin, 150mM NaCl, 50mM Tris pH 7.4, 2mM MgAc, and 1x Protease inhibitor cocktail) for 30min on ice. The cell lysates were then diluted with 600μL Buffer B (0.1% digitonin, 150mM NaCl, 50mM Tris pH 7.4) and cleared by centrifugation at 20,000g for 15 min. The supernatants were incubated with either anti-FLAG beads (Biolegend) or anti-HA magnetic beads (Thermo Scientific) for 1.5 hours. The beads were then washed for 3 times with 1 ml of Buffer B. The bound material was eluted from the beads with 50μL 2x SDS sample buffer and boiled for 5 min. The samples were then analyzed by immunoblotting with the indicated antibodies.

To test the interaction between WRB variants and Sec63 in Figure 5E, Sec63−/− cells stably expressing Sec63-FLAG (0.6 × 10^6^ /well) and HEK 293-Flp-In T-Rex cells were plated on poly-lysine coated 6-well plates and transiently transfected with either 2μg of WRB-HA-NFT variants or empty vector. After 24 hours of transfection and induction with 200ng/mL doxycycline, the aforementioned digitonin-based immunoprecipitation protocol was used and analyzed by immunoblotting with anti-Sec63, anti-Sec61α and anti-HA antibodies.

To test the interaction between WRB-NFT-Venus variants and the endogenous Hrd1 in Figure 6G, HEK 293-Flp-In T-Rex cells (0.6 × 10^6^ /well) were plated on poly-lysine coated 6-well plates and transiently transfected with either 1ug of pcDNA/FRT/TO-WRB-NFT-Venus variants or empty vector. After 24 hours transfection and induction with doxycycline, cells were treated with 10μM P97 inhibitor for 2 hours and harvested in PBS. The aforementioned digitonin-based immunoprecipitation protocol was used but with incubating the lysates with anti-GFP antibody for 1.5 hours followed by incubating with protein A agarose for another 1 hour. The samples were then analyzed by immunoblotting with anti-Hrd1, anti-Sel1 and anti-GFP antibodies.

To examine the interaction between BiP and WRB variants in Figure 6H, HEK 293-Flp-In T-Rex cells (0.6 × 10^6^ /well) were plated on poly-lysine coated 6 well plates and transiently transfected with either 1ug pcDNA/FRT/TO-WRB variants or empty vector. After 24 hours transfection and induction with doxycycline, the cells were harvested in PBS and centrifuged for 2 min at 10,000g. The cell pellets were lysed in NP40 buffer (50 mM Tris, pH 7.5, 150 mM NaCl, 0.5% deoxycholic acid, and 0.5% NP40) containing apyrase (10 U/ml), 10 mM CaCl_2_ and 1X protease inhibitor cocktail for 30 min on ice. The cell lysate was centrifuged at 15,000g for 15 min. The supernatant was incubated with anti-HA magnetic beads (Thermo Scientific) for 2 hours in the cold room. The beads were washed 3 times with 1 ml of NP40 buffer and eluted by directly boiling in 50 μl of 2x SDS sample buffer for 5 min and analyzed by immunoblotting with anti HA and anti-BiP antibodies.

### Purification of WRB and *in vitro* pull-down assay

To test the interaction between CAML and WRB variants *in vitro*, 293T cells (3 × 10^6^ /plate) were seeded on two poly-lysine coated 10cm plates and transfected with 12μg plasmid encoding either WRB-FLAG or WRB-K164L-FLAG. After 24 hours transfection, cells were trypsinized and transferred into 2× 15cm plates. After 48 hours transfection, cells were harvested and solubilized with 2mL lysis buffer (50mM Tris pH 8.0, 300mM NaCl, 5mM EDTA, 1% Triton X-100, 1× Protease inhibitor cocktail) for 30 min at 4°C. The lysates were then centrifuged at 20,000g for 30 min. The supernatant was incubated with anti-FLAG beads (Sigma) at 4°C for 2 hours in a 2ml Bio-Rad column. The beads were washed for 4 times with 5ml of wash buffer 1 (50mM Tris pH 8.0, 300mM NaCl, 5mM EDTA, 0.1% Triton X-100) and then washed 3 times with 2mL of wash buffer 2 (50mM Hepes pH 7.4, 100mM KAC, 50mM NaCl, 10mM MgAC, 0.2% DBC, 1mM DTT) to exchange Triton X-100 to DBC. The bound material was eluted from the beads with 300μL elution buffer (1mg/mL FLAG peptides in wash buffer 2). The relative concentration of WRB-FLAG and WRB (K164L)-FLAG were determined by immunoblotting. For the CAML pull down assay, 293T cells (0.6 × 10^6^ /well) were plated on poly-lysine coated 6-well plates and transfected with 2μg pcDNA/FRT/TO-CAML-HA. After 24 hours transfection, cells were harvested in PBS and centrifuged for 2 min at 10,000g. The cell pellets were lysed with 200μL lysis buffer 2 (50mM Tris pH7.4, 300mM NaCl, 2mM MgAC, 1% Triton X-100, 1X protease inhibitor cocktail) for 30 min on ice. The cell lysates were then diluted with 600μL wash buffer 3 (50mM Tris pH7.4, 300mM NaCl, 2mM MgAC, 0.1% Triton X-100) and cleared by centrifugation at 20,000g for 15 min. The supernatants were incubated with anti-HA magnet beads (Thermo Scientific) for 1.5 hours. The beads were then washed for 3 times with 1 ml of wash buffer 3 followed by washing with 1mL of wash buffer 2 for two times. Equal amount of purified WRB-FLAG or WRB (K164L)-FLAG was incubated with CAML bound anti-HA magnetic beads for 30min at 4°C. The beads were washed twice with 1mL of wash buffer 2, and the bound materials were eluted with 50μL 2× SDS buffer and analyzed by immunoblotting with anti-HA and anti-FLAG antibodies.

### Immunofluorescence

U2OS cells (0.1 × 10^6^/well) were plated on 12 mm round glass coverslips (Fisher Scientific) coated with 0.15mg/mL poly-lysine for 1 hour in 24-well plates and transfected with 400ng plasmids encoding WRB-HA or WRB-NFT-Venus using 1ul of Lipofectamine 2000. After 24 hours of transfection, cells were fixed with 3.7% formaldeyhyde (J.T. Baker, Phillipsburg, New Jersey) for 10 min and permeabilized with 0.1% Triton X-100 (American Analytical, Akron, OH) for 5 min. The non-specific binding sites were blocked with Buffer PSP (1xPBS containing 10% Horse Serum and 0.1% Saponin) for 45 min. 150μL of rabbit anti-HA, or rabbit anti-GFP primary antibodies were added at 1:100 dilution in Buffer PSP and incubated for 1 hour followed by washing for 5 times with each time incubating with buffer for 5 min. 150μL of the secondary antibodies anti-rabbit Cy3 (Jackson Immuno Research) were added at 1:100 dilution in Buffer PSP and incubated for 1 hour before washing five times with Buffer PSP. Coverslips were then incubated with 5 mg/ml Hoechst stain in 1xPBS for 15 min, washed with 1×PBS, and mounted using Fluoromount G (SouthernBiotech). Cells were imaged on Leica scanning confocal microscopy consisting of an inverted microscope (Leica SP6/SP8), and an HC PL APO 63X (CS2 No: 11506350) oil objective lens (Leica, Wetzlar, Germany), and was controlled by the Leica Application Suite X. Sequential image scanning at 3× zoom, 100 Hz, 1024 _1024 pixels, and with line averaging set at four was used to collect images.

### Quantification and statistical analysis

Quantification of western blot and autoradiographs were performed with image J gel analysis/ lane plotting. Error bars reflect the three independent experiments standard deviation.

## Supplementary Information

**Figure S1 (related to Figure 1–3).**
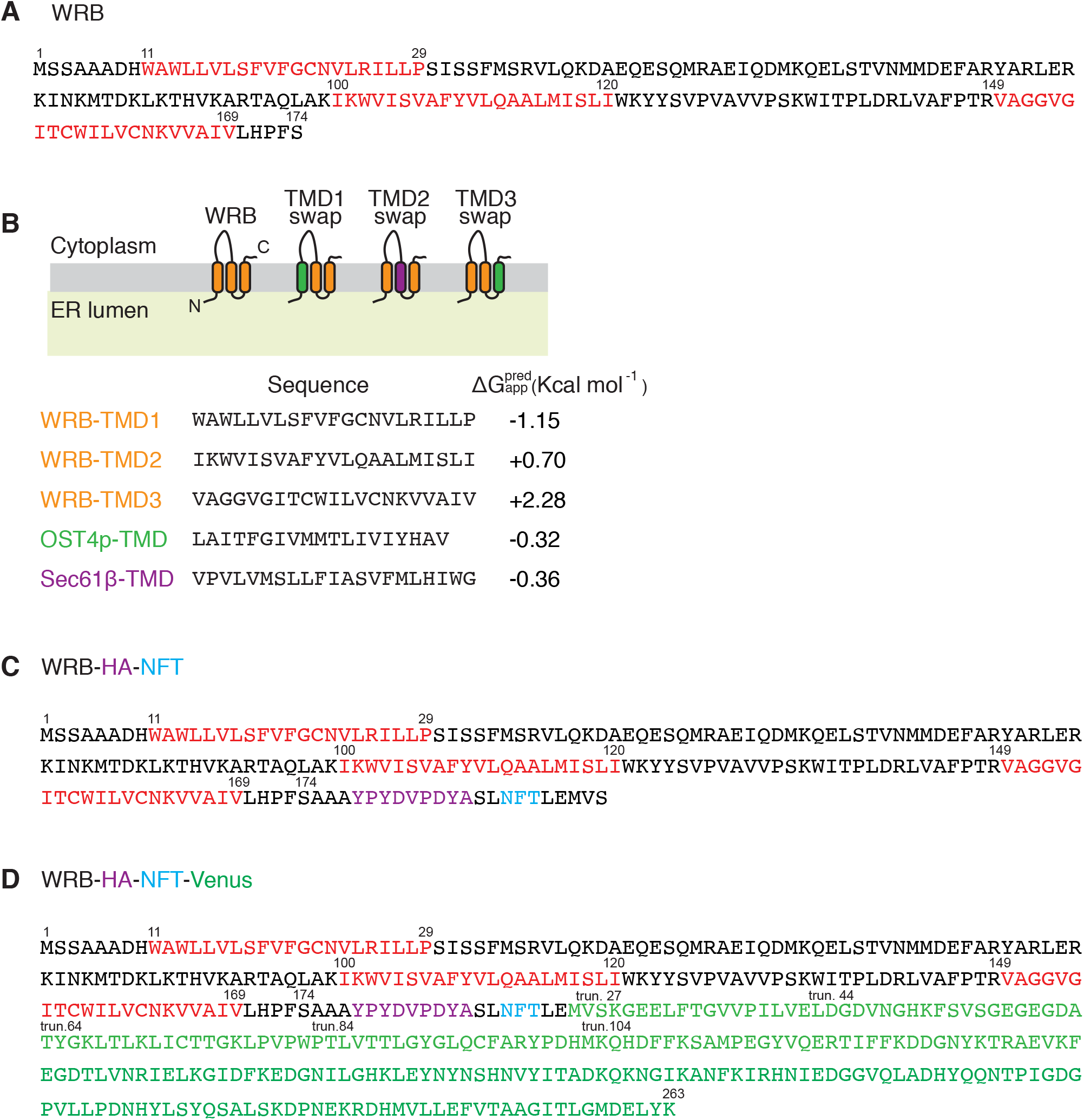
The amino acid sequences of WRB constructs. **(A)** The amino acid sequence of wild type WRB. The three transmembrane domains of WRB are colored in red. **(b**) The transmembrane domain sequences of WRB TMD1, TMD2, TMD3, OST1p, and Sec61β, the latter two were for swapping WRB TMDs. The apparent free energy values were listed next to the sequences. **(C)** The amino acid sequence of WRB-HA-NFT construct. **(D)** The amino acid sequence of WRB-HA-NFT-Venus construct.

**Figure S2 (related to Figure 2).**
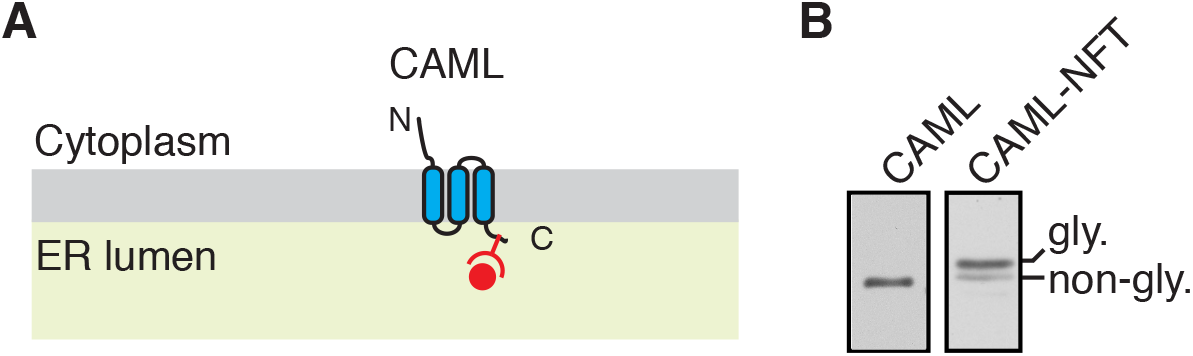
Probing CAML topology by glycosylation. **(A)** A schematic showing the predicted topology with an N-glycosylation motif (NFT) fused to the C-terminus of CAML. **(B)** CAML-HA and CAML-HA-NFT expressing cells were analyzed by immunoblotting with anti-HA antibody. Note that most of CAML-HA-NFT is glycosylated, suggesting that it is correctly oriented in the membrane.

**Figure S3 (related to Figure 2).**
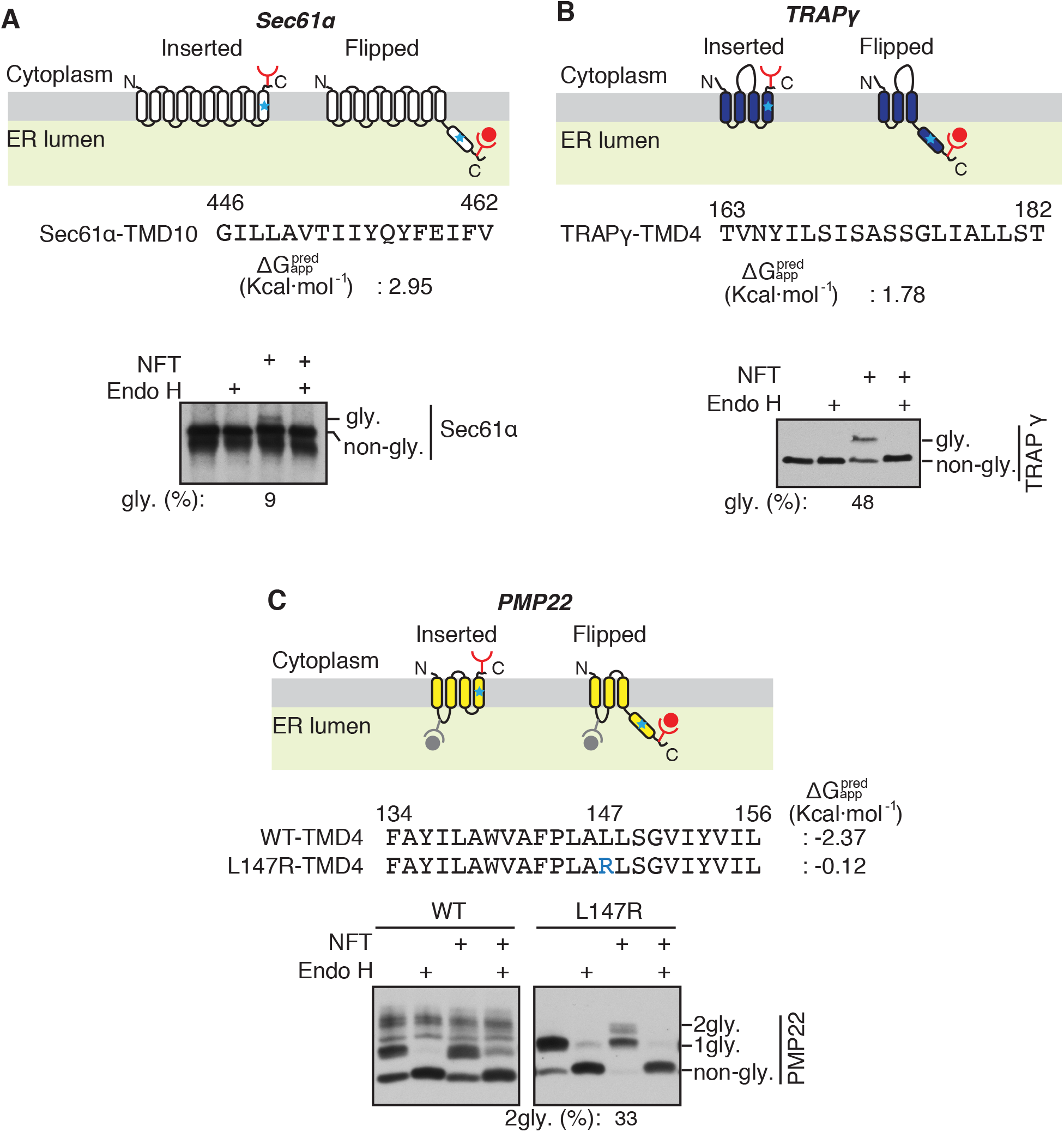
C-TMDs of unassembled membrane proteins are flipped into the ER lumen. **(A) T**op panel: A diagram showing the topology of Sec61α with both inserted and flipped forms. The amino acid sequence of C-TMD (TMD10) of dog Sec61α-TMD10 and its apparent free energy value are shown. Bottom panel: The lysates from either Sec61α-HA or Sec61α-HA-NFT transfected cells were treated without or with endoglycosidase H (Endo H) and analyzed by immunoblotting with anti-HA antibody. The percentage of flipped form (glycosylated) from total was quantified with Image J and shown under the immunoblot. **(B)** TRAPγ-HA-NFT and TRAPγ-HA were analyzed as in **A**. **(C)** PMP22-HA and PMP22 (L147R)-HA with and without the C-terminus glycosylation tag NFT were analyzed as in **A**. “2gly.” indicates the flipped form.

**Figure S4 (related to Figure 3).**
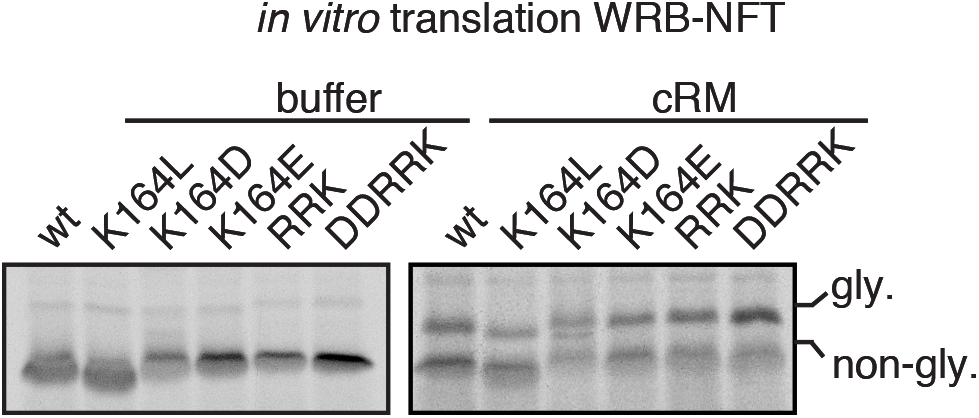
In vitro flipping of WRB constructs with short C-terminal tails. The indicated WRB-HA-NFT variants were in vitro translated in the absence or presence of canine pancreas rough microsomes (cRM). The translated products were analyzed by SDS-PAGE and autoradiography.

**Figure S5 (related to Figure 5).**
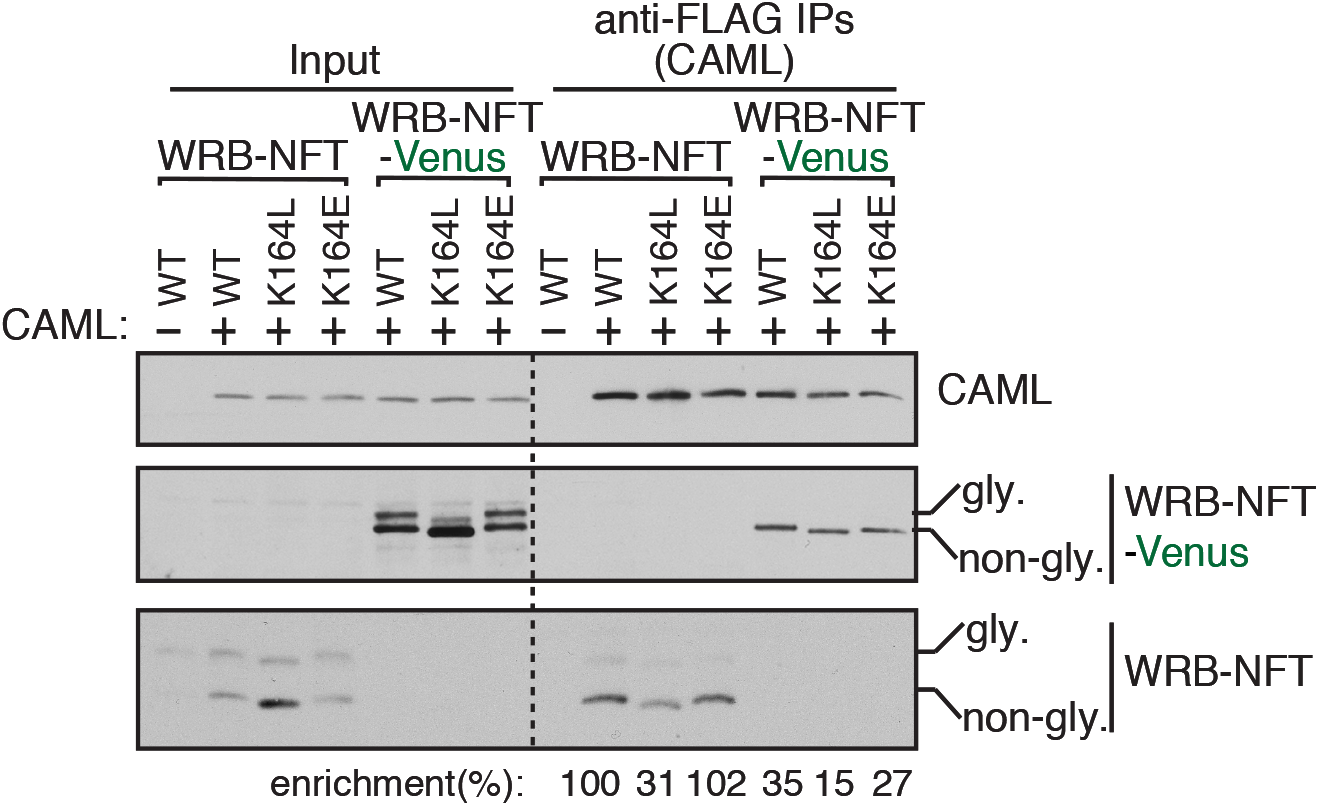
The C-TMD with a long tail inefficiently assembles with the partner TMD. The indicated WRB-HA-NFT or WRB-HA-NFT-Venus variants were co-transfected with CAML-FLAG into HEK293 cells and immunoprecipitated with anti-FLAG beads. The resulting samples were analyzed by immunoblotting with anti-FLAG antibody (CAML) and anti-HA antibody (WRB-HA-NFT and WRB-HA-NFT-Venus variants). The band intensity was quantified by Image J, and the ratios of IPs to inputs were calculated. The IP enrichment of WRB-NFT wild type was taken as 100%.

**Figure S6 (related to Figure 5).**
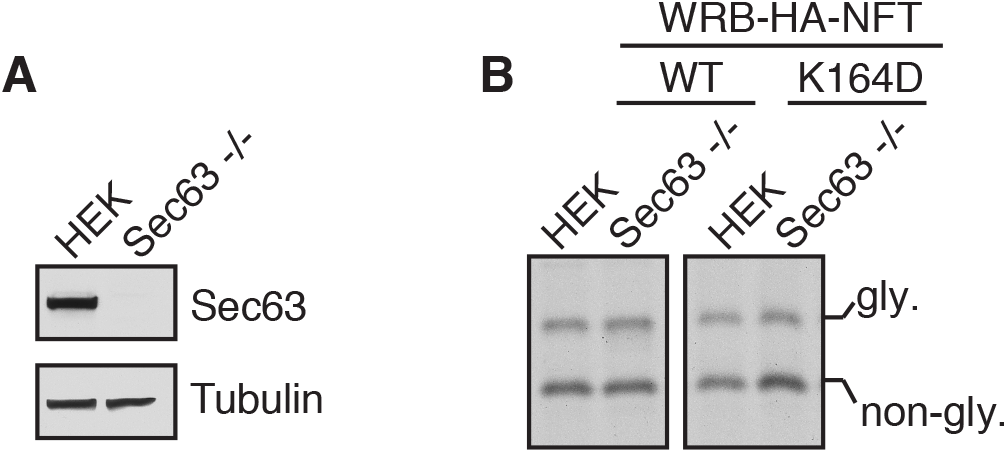
Sec63 is not required for the flipping of C-TMD. **(A)** The lysate of HEK293 or Sec63−/−cells was immunoblotted for the indicated antigens. **(B)** The indicated WRB-HA-NFT constructs were transfected into HEK293 and Sec63−/−cells. The cells were metabolically labeled and analyzed by autoradiography after immunoprecipitation with anti-HA beads.

**Figure S7 (related to Figure 6).**
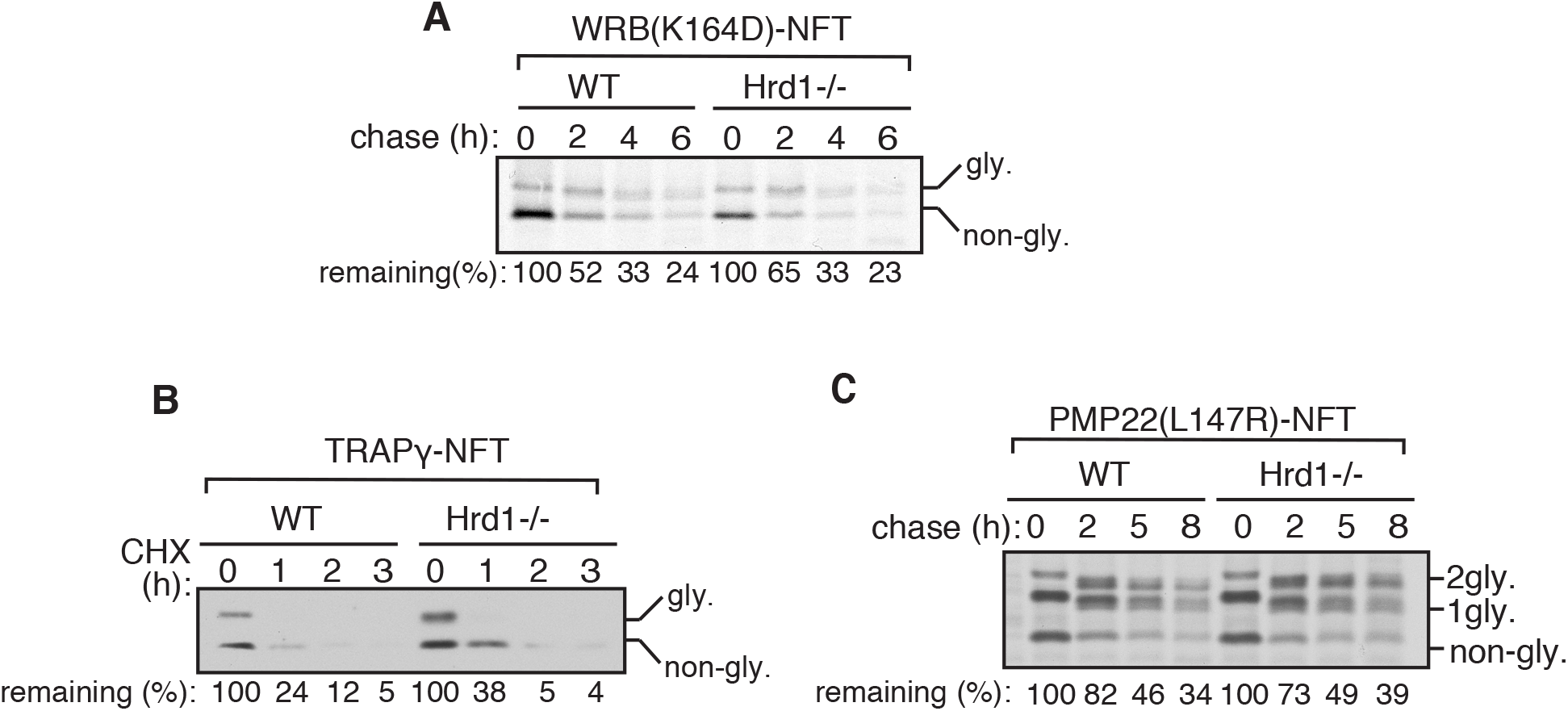
Hrd1 independent degradation of flipped C-TMDs with short cytosolic tails. **(A)** HEK293 or Hrd1 −/− cells expressing WRB (K164D)-HA-NFT were metabolically labeled and chased for the indicated time points and analyzed by autoradiography after immunoprecipitation with anti-HA beads. The protein level at 0-hour time point was taken as 100%, and the percentage of the remaining protein was calculated with respect to 0 hour. **(B)** TRAPγ-HA-NFT was transfected into either HEK293 or Hrd1−/− cells. After 24h of transfection, cells were treated with cycloheximide (CHX) for the indicated time points and analyzed by immunoblotting for the indicated antigens. The percentage of remaining protein level was calculated as in **A.** **(C)** PMP22(L147R)-NFT was transfected into either HEK293 or Hrd1−/− cells and analyzed as in **A.**

**Figure S8 (related to Figure 6).**
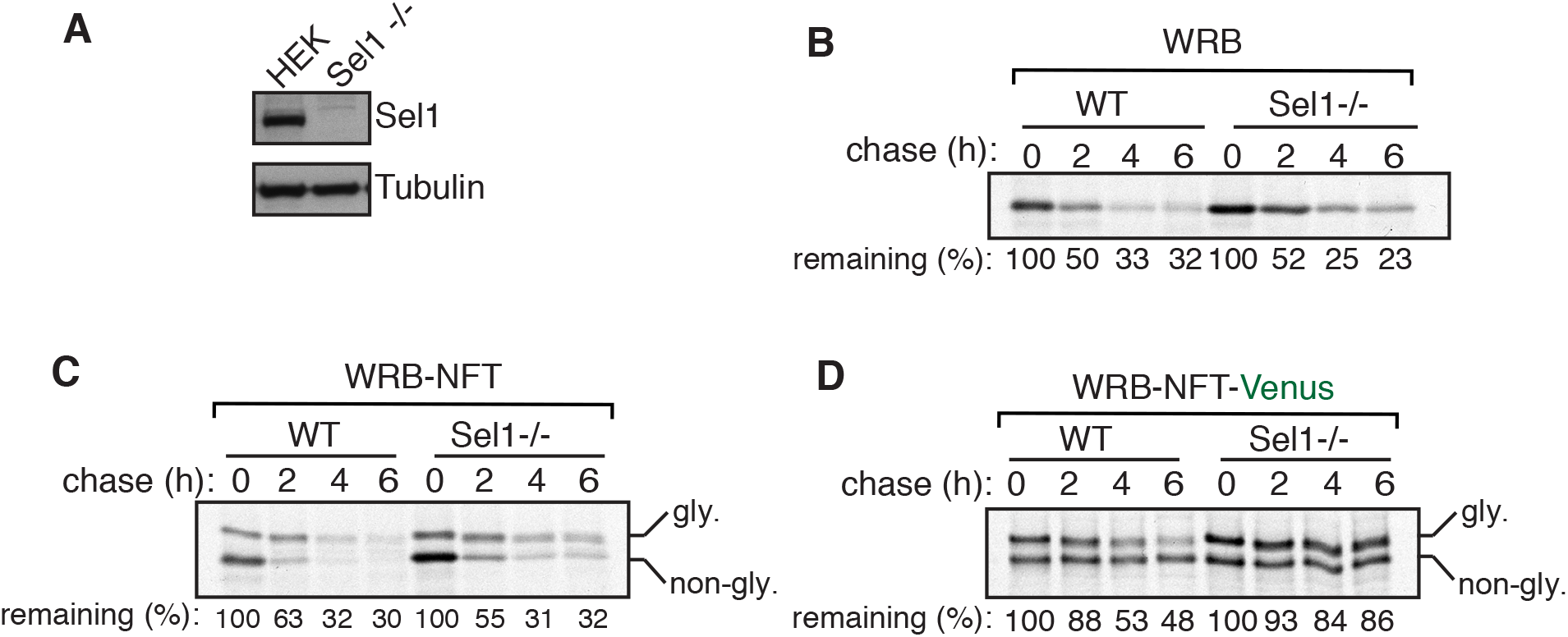
Sel1 dependent or independent degradation of flipped C-TMD substrates. **(A)** The lysate of HEK293 or Sel1−/− cells was immunoblotted for the indicated antigens. **(B)** HEK293 or Sel1−/− cells expressing WRB-HA were metabolically labeled and chased for the indicated time points. The samples were analyzed by autoradiography after immunoprecipitation with anti-HA beads. The protein level at 0-hour time point was taken as 100%, and the percentage of the remaining protein was calculated with respect to 0 hour. **(C-D)** The indicated WRB variants were transfected into HEK293 or Hrd1−/− cells and analyzed as in **B.** For WRB-NFT-Venus, immunoprecipitation was conducted with anti-GFP antibodies.

